# Depletion of limiting rDNA structural complexes 1 triggers chromosomal instability and replicative aging of *Saccharomyces cerevisiae*

**DOI:** 10.1101/380675

**Authors:** Ryan D. Fine, Nazif Maqani, Mingguang Li, Elizabeth Franck, Jeffrey S. Smith

**Affiliations:** Department of Biochemistry and Molecular Genetics, University of Virginia School of Medicine, Charlottesville, VA 22908; Department of Laboratory Medicine, Jilin Medical University, Jilin, 132013, China

**Keywords:** Sir2, cohesin, RENT, monopolin, yeast, replicative lifespan, aging, rDNA

## Abstract

Sir2 is a highly conserved NAD^+^-dependent histone deacetylase that functions in heterochromatin formation and promotes replicative lifespan (RLS) in the budding yeast, *Saccharomyces cerevisiae*. Within the yeast rDNA locus, Sir2 is required for efficient cohesin recruitment and maintaining stability of the tandem array. In addition to the previously reported depletion of Sir2 in replicatively aged cells, we discovered that subunits of the Sir2 containing complexes, SIR and RENT, were depleted. Several other rDNA structural protein complexes also exhibited age-related depletion, most notably the cohesin complex. We hypothesized that mitotic chromosome instability (CIN) due to cohesin depletion could be a driver of replicative aging. ChIP assays of the residual cohesin (Mcd1-13xMyc) in moderately aged cells showed strong depletion from the rDNA and initial redistribution to the point centromeres, which was then lost in older cells. Despite the shift in cohesin distribution, sister chromatid cohesion was partially attenuated in aged cells and the frequency of chromosome loss was increased. This age-induced CIN was exacerbated in strains lacking Sir2 and its paralog, Hst1, but suppressed in strains that stabilize the rDNA array due to deletion of *FOB1* or through caloric restriction (CR). Furthermore, ectopic expression of *MCD1* from a doxycycline-inducible promoter was sufficient to suppress rDNA instability in aged cells and to extend RLS. Taken together we conclude that age-induced depletion of cohesin and multiple other nucleolar chromatin factors destabilize the rDNA locus, which then results in general CIN and aneuploidy that shortens RLS.

## Introduction

Budding yeast replicative lifespan (RLS) was originally described decades ago as the number of times a mother cell divides before losing viability (Mortimer and Johnston 1959), and has been an effective model system for the identification and/or characterization of several conserved aging-related genes and pathways, including *SIR2*, AMPK (Snf1), and TOR signaling (Wasko and Kaeberlein 2014). *SIR2* is possibly the most famous yeast gene associated with replicative aging and encodes the founding family member of the NAD^+^-dependent histone/protein deacetylases, commonly known as sirtuins (reviewed in (Buck *et al*. 2004)). The NAD^+^ dependence of sirtuins provides a direct link between metabolism and cellular processes regulated by these enzymes. In fact, recent evidence points to depletion of cellular NAD^+^ pools as a potential mechanism for aging-associated disease, which could be mediated by impairment of sirtuins or other NAD^+^ consuming enzymes (Gomes *et al*. 2013). Therefore, understanding how sirtuins are impacted by aging and how they regulate age-altered cellular processes is of intense interest.

Eukaryotic genomes generally encode for several sirtuin homologs. The *Saccharomyces cerevisiae* genome, for example, encodes *SIR2* and four additional <u>H</u>omologs of <u>S</u>ir <u>T</u>wo (*HST1-HST4*) (Brachmann *et al*. 1995; Derbyshire *et al*. 1996). Sir2 and its fellow <u>S</u>ilent <u>I</u>nformation <u>R</u>egulator proteins, Sir1, Sir3, and Sir4 were originally shown to establish and maintain silencing of the silent mating loci, *HML* and *HMR* (Rine and Herskowitz 1987). These proteins form the so-called SIR complex that is recruited to and then spreads across the *HM* loci and telomeres to form hypoacetylated heterochromatin-like domains (reviewed in (Gartenberg and Smith 2016)). Sir2 is required for replicative longevity and its abundance is significantly reduced in replicatively aged yeast cells (Dang *et al*. 2009), presenting a possible mechanism for the decline of Sir2-dependent processes during aging. including gene silencing. Indeed, the depletion of Sir2 in aged cells causes hyperacetylated H4K16 and silencing defects at subtelomeric loci (Dang *et al*. 2009). It has been reported that aged cells become sterile (mating-incompetent) due to loss of silencing at *HML* and *HMR* (Smeal *et al*. 1996), which results in co-expression of the normally repressed α1/α2 and a1/a2 transcription factor genes encoded at these loci. In theory, this should induce a diploid-like or “pseudodiploid” gene expression pattern and sterility, as is observed for a silencing defective *sir2*Δ mutant (Rine and Herskowitz 1987). However, more recent experiments point toward a silencing-independent mechanism of sterility, whereby aggregation of the Whi3 protein in aged cells makes them insensitive to pheromones (Schlissel *et al*. 2017).

Alternative models for Sir2 control of RLS have focused on the rDNA tandem array where Sir2 is important for cohesin recruitment (Kobayashi *et al*. 2004; Ganley and Kobayashi 2014). Cohesin association with the rDNA also requires Tof2 and the Lrs4/Csm1 (cohibin) complex (Huang *et al*. 2006). Sir2 silences RNA polymerase II-dependent transcription at the rDNA locus via a nucleolar HDAC complex called RENT (Bryk *et al*. 1997; Smith and Boeke 1997), consisting of Sir2, Net1, and Cdc14 subunits (Shou *et al*. 1999; Straight *et al*. 1999). Specifically, RENT represses transcription of endogenous non-coding RNAs from the intergenic spacer (IGS) regions (Li *et al*. 2006). Derepression of the bidirectional promoter (E-pro) within IGS1 in *sir2*Δ cells displaces cohesin from the rDNA, thus destabilizing the array by making it more susceptible to unequal sister chromatid exchange (Kobayashi and Ganley 2005). Mild Sir2 overexpression, on the other hand, enhances silencing, suppresses recombination between repeats, and extends RLS (Smith *et al*. 1998; Kaeberlein *et al*. 1999).

Extrachromosomal rDNA circles (ERCs) derived from these unequal recombination events specifically accumulate to high levels in old mother cells (Sinclair and Guarente 1997), where they can interfere with G1 cyclin expression (Neurohr *et al*. 2018). Such an ERC-centric model is supported by RLS extension of *fob1*Δ strains (Defossez *et al*. 1999). Fob1 binds to the rDNA at IGS1 to block DNA replication forks from colliding with elongating RNA polymerase I molecules (Kobayashi and Horiuchi 1996). The blocked forks can collapse, resulting in double-stranded DNA breaks that trigger unequal sister chromatid exchange (Takeuchi *et al*. 2003). The frequency of rDNA recombination and ERC production is reduced in a *fob1*Δ mutant due to loss of the fork block, thus extending RLS (Defossez *et al*. 1999). More recently, this rDNA-centric model of aging has been extended to include general rDNA instability having negative effects on genome integrity, including ERC accumulation, and is also considered a critical contributor to aging (Ganley and Kobayashi 2014).

In addition to promoting cohesin recruitment to the rDNA, Sir2 is also required for establishing sister chromatid cohesion (SCC) at *HML* and *HMR* (Chang *et al*. 2005; Wu *et al*. 2011). Moreover, we previously observed significant overlap between Sir2 and cohesin at additional binding sites throughout the genome (Li *et al*. 2013). Outside heterochromatin, the cohesin loading complex (Scc2/Scc4) deposits cohesin (Mcd1, Irr1, Smc1, Smc3) onto centromeres and other cohesion associated regions (CARs) (Ciosk *et al*. 2000; Kogut *et al*. 2009), in order to maintain SCC until anaphase, when Mcd1 is cleaved by separase to facilitate sister chromatid separation (reviewed in (Marston 2014)). Cohesin defects therefore result in chromosome instability (CIN) due to improper chromosome segregation (reviewed in (Wood *et al*. 2010)). A previous study found that cells deleted for *SIR2* have a CIN phenotype related to hyperacetylation of H4K16 (Choy *et al*. 2011), though the functional relationship to cohesin was not considered. Given the natural depletion of Sir2 from replicatively aging yeast cells (Dang *et al*. 2009), we hypothesized the frequency of CIN should increase with age. Here, we establish that CIN is indeed more frequent in aged cells and is associated with SCC defects. We go on to show that like Sir2, cohesin subunits are depleted from aged mother cells, providing a likely reason for problems with SCC. Interestingly, despite the overall reduction in cohesin protein levels, the chromosomal distribution of cohesin enrichment was not uniform. In moderately aged mother cells, enrichment at the rDNA was drastically reduced, while binding at centromeres was not, suggesting a mechanism by which SCC is preferentially maintained at centromeres to ensure cell viability. However, this comes at the expense of chronic rDNA instability that is exacerbated by additional age-induced reductions in the RENT and cohibin/monopolin complexes. The defects in rDNA stability can be suppressed by overexpressing the Mcd1 subunit of cohesion, which also extends RLS, thus making cohesin a dose dependent longevity factor. Lastly, the age associated CIN phenotype is suppressed by deleting *FOB1* or by CR growth conditions, suggesting a model whereby rDNA instability on chromosome XII caused by RENT, cohibin, and cohesin depletion drives the mitotic segregation defects of other chromosomes during replicative aging.

## Methods

### Yeast strains, plasmids, and media

Yeast strains were grown at 30°C in Yeast Peptone Dextrose (YPD) or Synthetic Complete (SC) medium for strains bearing plasmids (Matecic *et al*. 2010). *SIR2, HST1*, or *FOB1* open reading frames were disrupted with one-step PCR-mediated gene replacement using *kanMX4*, *natMX4*, or *hphMX4* drug resistance markers, respectively. The *HMR* deletion by replacement with *hphMX4* spans sacCer3 genome chrIII coordinates 293170-294330. All C-terminally 13xMyc (EQKLISEEDL) tagged proteins were targeted at endogenous loci in the Mother Enrichment Program (MEP) strains UCC5181 and UCC5179 (Lindstrom and Gottschling 2009), followed by mating to generate homozygous diploids. All deletions and fusions were confirmed by colony PCR, western blotting, or both. pRF4 was constructed by PCR amplifying the *MCD1* open reading frame from ML1 genomic DNA and ligating into *Pst*I and *Not*I sites of pCM252 (Belli *et al*. 1998), a tetracycline inducible overexpression vector (Euroscarf). pRF10 and pRF11 were constructed by removing the expression cassette by *Pvu*II blunt end digestion of pCM252 and pRF4, respectively, and ligating it between the *Pvu*II sites of pRS405 bearing *LEU2*, thus replacing the *TRP1* marker with *LEU2*. pRF10 and 11 were then digested with *EcoRV* and integrated into the genome at *leu2Δ1*. pSB760 and pSB766 are integrating and *2μ LEU2* vectors, respectively, bearing a single copy of *SIR2* (Buck *et al*. 2002). All strains used in this work are listed in Table S1 and all primers are listed in Table S2.

### Isolation of aged yeast cells

Aged yeast cell enrichment was based on the MEP strain background (Lindstrom and Gottschling 2009; Lindstrom *et al*. 2011). For all assays, 1 μL of stationary phase culture was inoculated into 100 mL of YPD medium and then grown into log phase. Approximately 1×10^8^ cells were harvested, and centrifuged cell pellets washed 3 times with 1x phosphate buffered saline (PBS). Cells were then resuspended in 1 mL PBS and mixed with 5 mg of Sulfo-NHS-LC-Biotin (Pierce) per 1×10^8^ cells for 30 min at room temperature. After biotin labeling, 5×10^7^ cells were added to 1.5 L YPD cultures containing 1 μM estradiol, and 100 μg/mL ampicillin to prevent bacterial contamination. These cultures were allowed to grow for 24 hr to 36 hr before processing in an assay specific manner (see below). For non-MEP strain backgrounds, estradiol was not added to the cultures.

### Western blotting

Two 1.5 L cultures were used for each western blot experiment, corresponding to approximately 2×10^7^ total aged cells after purification for each biological replicate. Cells were pelleted using a Sorvall RC-5B Plus centrifuge with an SLA-3000 rotor at 2000 rpm, then resuspended at a density of 6×10^8^ cells/mL in RNAlater (Ambion) for 45 min in two separate conical tubes. Following fixation, cells were pelleted and resuspended in 45 mL of cold PBS, 2 mM EDTA in 50 mL conical tubes. The mixture was incubated at 4°C for 30 min with 800 μL of Streptavidin MicroBeads (Miltenyi Biotec), which were then purified through an autoMACS Pro Cell Separator using the posseld2 program (UVA Flow Cytometry Core Facility). A 20 μL aliquot of each output was used for bud scar counting using calcofluor white staining, before combining the isolated samples into a single microfuge tube. Samples were frozen at −80°C before protein extraction. Thawed cells were vortexed twice for 1 min in 20% TCA (trichloroacetic acid) with ∼100 μL of acid washed glass beads with a brief cooling period in between vortexing. Beads were allowed to settle before transferring the supernatant to a fresh microfuge tube. A 250 μL wash of 5% TCA was applied twice to the beads and pooled with the initial lysis sample. Proteins were precipitated at 10,000 rpm in a microfuge for 5 min at 4°C. Pellets were then resuspended in 50 μL of 1x SDS sample buffer (50 mM Tris-HCl pH 6.8, 2% SDS, 10% Glycerol, 3.6 M 2-mercaptoethanol) and neutralized with 30 μL of 1M Tris-HCl, pH 8.0. Samples were run on a 9% (w/v) SDS-polyacrylamide gel and transferred to an Immobilon-P membrane (Millipore). Membranes were incubated for 1 hour at room temperature in 1xTBST + 5% non-fat milk with primary antibodies (1:2000 α-Myc 9E10, 1:5000; α-Vma2 (Life Technologies); 1:5000 α-Sir2 (Santa Cruz Biotechnology); 1:1000 α-Sir4 (Santa Cruz Biotechnology); 1:1000 α-Sir3 (Santa Cruz Biotechnology). HRP-conjugated secondary antibodies (Promega) were diluted 1:5000, and detected using chemiluminescence with HyGLO (Denville Scientific). Quantitation was performed with ImageJ by using the rectangle tool to outline protein bands and an equivalent sized box for background. After subtraction of background, the signal of the aged cell band was divided by the signal of the Vma2 loading control, and finally normalized to the quantity of the Vma2 normalized young cell band which was arbitrarily set at 1.0.

### ChIP Assays with aged and young cell populations

Two 1.5 L cultures were used for each biological replicate. After centrifugation, cells were washed with 1xPBS then resuspended with 45 mL of PBS and incubated with 800 μL of streptavidin microbeads, followed by sorting with the autoMACS Pro Cell Separator. Sorted cells were immediately crosslinked with 1% formaldehyde for 20 min at room temperature, then transferred to screw cap microcentrifuge tubes and the pellets flash frozen in liquid nitrogen. Cells were thawed and lysed in 600 μL FA140 Lysis buffer (50 mM HEPES, 140 mM NaCl, 1% Triton X-100, 1 mM EDTA, 0.1% SDS, 0.1 mM PMSF, 1x protease inhibitor cocktail; Sigma) by shaking with acid-washed glass beads in a Mini-Bead beater (Biospec Products). Cell lysates were recovered and sonicated for 30 cycles of 30 sec “on” and 30 sec “off” in a Diagenode Bioruptor followed by centrifugation at 16,000 x g. A 1/10^th^ supernatant volume input was taken for each sample and crosslinking reversed by incubating overnight at 65°C in 150 μL elution buffer (TE, 1% SDS). The remaining supernatant was used for immunoprecipitation overnight at 4°C with 5 μg of primary antibody and 30 μL of protein G magnetic beads (Pierce), followed by washing 1x with FA-140 buffer, 2x with FA-500 buffer (FA-140 with 500 mM NaCl), and 2x with LiCl solution (10 mM Tris-HCl, pH 8.0, 250 mM LiCl, 0.5% NP-40, 0.5% SDS, 1 mM EDTA). DNA was eluted twice with 75 μL of elution buffer in a 65°C water bath for 15 min. The eluates were combined and crosslinking reversed. Input and ChIP DNA samples were purified by an Invitrogen PureLink™ PCR purification kit. Finally, ChIP DNA was quantified by real-time PCR and normalized to the input DNA signal. Young cells were collected flow-through from the autoMACS cell sorter and then processed as described for the aged cells.

### Sister chromatid cohesion assay

From 50 mL log phase SC cultures of strains 3349-1B, 3312-7A, 3460-2A, RF258, RF278, and RF290, 5×10^7^ cells were washed and biotinylated as described in the Isolation of aged yeast cells section. This population was transferred into a 1.5 L SC culture and allowed to grow for 14 hr. The original biotinylated cells were then purified by incubation with 300 μL of streptavidin micro beads followed by gravity filtration through a Miltenyi LS column. The column was washed twice with 5 ml of PBS and then processed as described below for young cells.

From the original log phase culture, 5×10^7^ cells were transferred to a fresh 50 mL SC culture and arrested in mitosis with 10 μg/mL nocodazole for 1.5 hr. For the *mcd1-1* strain 3312-7A, cells were also shifted to 37°C at this time. For bud scar staining, 5 mg of calcofluor white was dissolved in 1 mL of PBS and any remaining aggregate removed by centrifugation. The 1 mL of soluble calcofluor was then added to the 50 mL SC culture. Non-arrested cells were directly stained with calcofluor. Cells were then pelleted and washed in PBS. Following staining, 200 μL of 4% paraformaldehyde was added directly to the cell pellet and allowed to crosslink for 15 min at room temperature. The cell pellet was washed once with PBS and resuspended in ∼100-200 μL of 0.1 MKPO_4_/1 M sorbitol, pH 6.5. Images were captured with a Zeiss Axio Observer z1 widefield microscope using a 64x oil objective lens.

### Replicative lifespan assays

Lifespan assays were carried out as previously described (Steffen *et al*. 2009). Briefly, small aliquots of log phase cultures were dripped in a straight line onto solid agar YPD with 2% glucose. From the initial populations, a minimum of 32 virgin daughter cells were picked for lifespan assays with daughter cells being selectively pulled away from mother cells using a fiberoptic dissection needle and on a Nikon Eclipse 400 microscope. All virgin daughters were required to bud at least one time to be included in the experiment and dissection was carried out over the course of several days with temporary incubation at 4°C in between dissection periods to stop division. Cells were considered dead when they stopped dividing for a minimum of 2 generation times (180 min). For p-values indicated in the text, a Wilcoxon rank-sum test was conducted for respective lifespan assays using the basic wilcox.test function in R.

### Mini-chromosome loss (sectoring) assay

The colony sectoring assay was performed on SC plates with adenine limited to 80 μM. Frequency of mini-chromosome loss represents the number of ½ red/white sectored colonies divided by the sum of sectored and white colonies. Cells were plated to an approximate density of 500 cells/plate based on counts from a Brightline hemocytometer. Any plates bearing greater than 1000 cells were discarded. Three biological replicates of each strain were performed, with at least 10 plates counted per replicate. For aged cell populations, ∼5×10^6^ biotinylated cells were aged in 1.5 L of YPD for 24 hr. Cells were incubated with 300 μL of streptavidin magnetic beads (New England Biolabs) and manually washed 4 times with PBS on a magnetic stand, then plated onto the limiting adenine SC plates such that ∼500 colonies appeared. Bud scars were not counted because the size of the beads prohibited visualization.

### RT-qPCR measurement of *MCD1* overexpression

Doxycycline was added to log phase cultures at a concentration of 2 μg/mL for 4 hr in order to induce expression of *MCD1*. Total RNA was extracted using a standard acid phenol extraction protocol (Ausubel *et al*. 2000). cDNA was created from 1 μg of RNA using a Verso cDNA synthesis kit (Thermo Fisher). *MCD1* expression levels were quantified on an Applied Biosystems StepOne real time PCR machine with primers JS2844 and JS2949, and normalized to actin transcript levels (primers JS1146 and JS1147).

### Hi-C library construction

Log-phase cultures were cross-linked with 3% formaldehyde for 20 min and quenched with a 2x volume of 2.5M glycine. Cell pellets were washed with dH_2_O and stored at −80°C. Thawed cells were resuspended in 5 ml of 1X NEB2 restriction enzyme buffer (New England Biolabs) and poured into a pre-chilled mortar containing liquid N_2_. Nitrogen grinding was performed twice as previously described (Belton and Dekker 2015), and the lysates were then diluted to an OD_600_ of 12 in 1x NEB2 buffer. 500 μl of cell lysate was used for each Hi-C library as follows. Lysates were solubilized by the addition of 50 μl 1% SDS and incubation at 65°C for 10 min. 55 μl of 10% Triton X-100 was added to quench the SDS, followed by 10 μl of 10X NEB2 buffer and 15 μl of *Hin*dIII (New England Biolabs, 20 U/μl) to digest at 37°C for 2 hr. An additional 10 μl of *Hin*dIII was added for digestion overnight. The remainder of the protocol was based on previously published work with minor exceptions (Burton *et al*. 2014). In short, ends were filled in with dNTPs and biotinylated dCTP at 0.4 mM concentration using Klenow Exo-(NEB) for 1 hr at 37°C. After a brief heat inactivation, blunt ends were ligated together in 3 mL reaction volumes with T4 DNA-ligase for 6 hr at 16°C with a minimum DNA concentration of 0.5 ng/μL. Following ligation, cross-links were reversed at 70°C O/N and DNA was purified by phenol/chloroform extraction and ethanol precipitation. Unligated-biotinylated ends were removed using T4 DNA Polymerase (NEB). DNA was purified one final time with two Zymogen DNA Clean and Concentrate-5 Kit columns per ligation reaction and eluted with 65 μL TE (130 μL total). Chromatin was quantitated with a Qubit fluorometer and then sheared using a Diagenode Bioruptor. Hi-C sequencing libraries were prepared with reagents from an Illumina Nextera Mate Pair Kit (FC-132-1001) using the standard Illumina protocol of End Repair, A-tailing, Adapter Ligation, and 12 cycles of PCR. PCR products were size selected and purified with AMPure XP beads before sequencing with an Illumina Miseq or Hiseq.

### Hi-C data analysis

Iteratively corrected heatmaps appearing in this manuscript were produced using python scripts from the Mirny lab hiclib library, publicly available at: http://mirnylab.bitbucket.org/hiclib/index.html. Briefly, reads were mapped using the iterative mapping program and then run through the fragment filtering program using the default parameters. Raw heat maps were further filtered to remove diagonal reads and iteratively corrected. Finally, the iteratively corrected heatmaps were normalized for read count differences to make them comparable. The *cdc15-2* sample data was pulled from the SRA database at SRP094582 (Lazar-Stefanita *et al*. 2017), and our data is available from GEO at GSE117037.

### Data availability

Strains and plasmids are available upon request. Supplemental Tables S1 and S2, containing lists of all strains and oligodeoxynucleotides used in the study, are included in the Supplemental Information file. Supplemental Figures S1 through S7 are also included in the same file.

## Results

### Sir2 binding partners are depleted in aged yeast cells

Sir2 is a dosage dependent longevity factor, such that strains provided one extra *SIR2* gene copy have an extended RLS (Kaeberlein *et al*. 1999). Mother cells experience a progressive reduction of Sir2 protein during normal replicative aging that presumably contributes to the aging process (Dang *et al*. 2009). Sir2 does not function in isolation, so we hypothesized that protein levels of key Sir2-interacting partners could also be depleted with age. To isolate sufficient quantities of aged cells for western blot assays we turned to <u>M</u>other <u>E</u>nrichment <u>P</u>rogram (MEP) strains developed by the Gottschling lab (Lindstrom and Gottschling 2009). The aged cell purification procedure was validated by increased average bud scar counts with calcofluor white and the expected reduction of Sir2 protein (Figure 1A). The vacuolar protein Vma2, used as a loading control for these western blots, does not deplete with age (Lindstrom *et al*. 2011). Since Sir2 is the catalytic subunit of both SIR and RENT (Figure 1B), it was important to know which complexes were impacted by age. As shown in Figure 1C and D, Sir4 was strongly depleted in aged cells while Sir3 was actually enriched. Sir3 enrichment in aged cells was also observed in an earlier proteomics screen (Janssens *et al*. 2015). Such a stark difference was considered relevant because Sir2 and Sir4 interact as a heterodimer that associates with the acetylated H4 N-terminal tail (Moazed *et al*. 1997), while Sir3 is subsequently recruited following H4K16 deacetylation to complete SIR holocomplex formation on heterochromatin (Oppikofer *et al*. 2011). Myc-tagged Net1 (RENT complex) was also strongly depleted from aged cells (Figure 1E), indicating that Sir2/Sir4 and the nucleolar RENT complex are both depleted during aging. It should be noted that the Sir2 paralog, Hst1, which has the capacity to compensate for loss of Sir2 (Hickman and Rusche 2007; Li *et al*.), was also partially depleted from aged cells (Figure 1F). Detailed bud scar and protein quantitations from triplicate biological samples are reported in Table S3.

**Figure 1.**
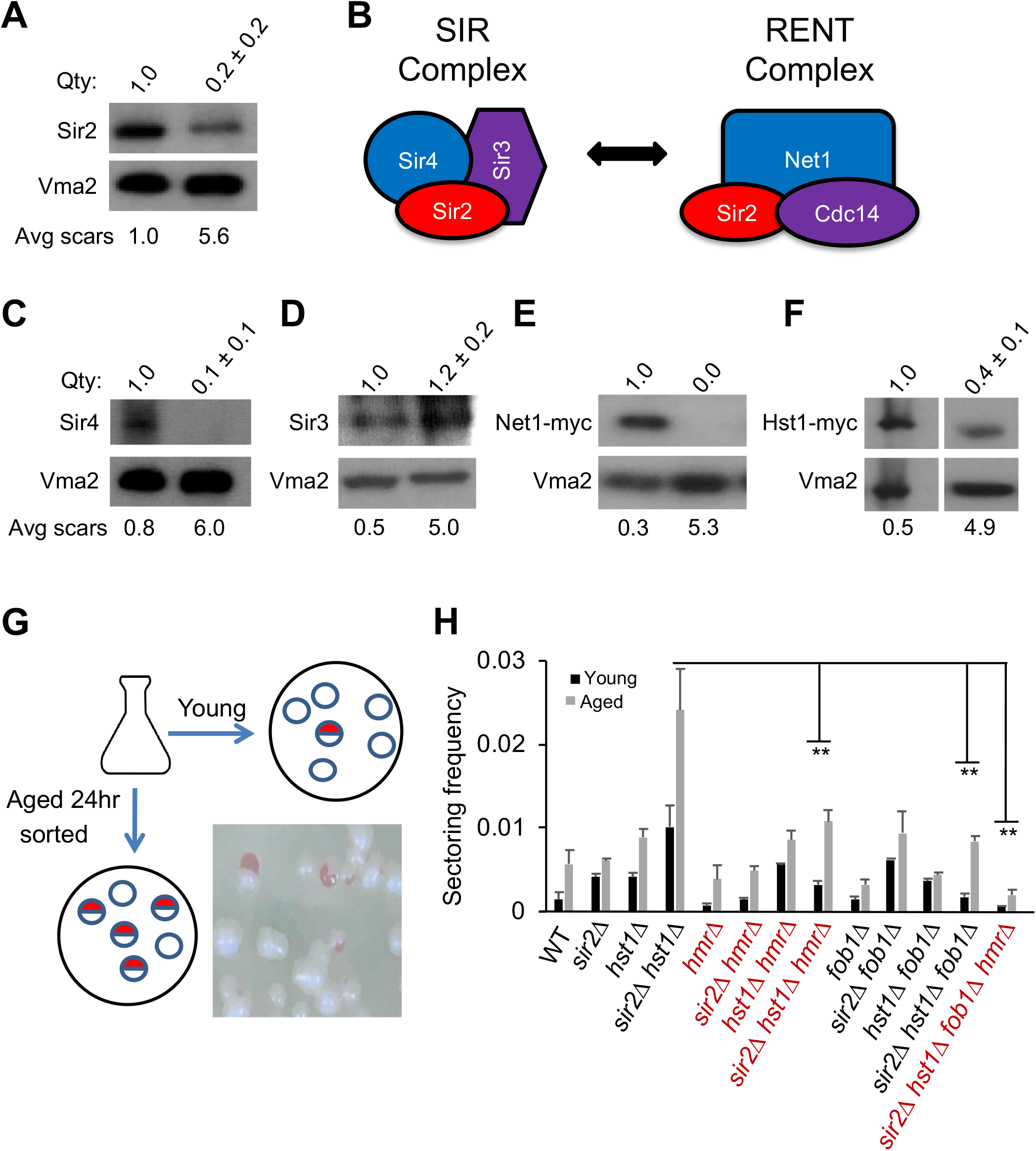
Depletion of Sir2 complexes and elevated chromosome instability in replicatively aging cells. **(A)** Western blot of Sir2 protein levels in young and aged cells. Vma2 serves as a loading control. **(B)** Depiction of the SIR and RENT complexes, which share Sir2 as a catalytic subunit. **(C, D)** Western blots of native Sir4 and Sir3. **(E, F)** Western blots of 13xMyc tagged Sir2 paralog, Hst1, and the Net1 subunit of RENT. **(G)** Schematic of artificial chromosome loss assay for ½ sectored colonies. **(H)** Quantification of sectoring frequency for young or aged cell populations. Approximately 10,000 colonies were analyzed for each strain across several biological replicates. *p<0.05, two-tailed student t-test. Bud scar counts are an average for each enriched population. Qty: indicates the mean western signal of each protein relative to Vma2, with the ratio for young cells set at 1.0 (n=3 biological replicates).

### Chromosome instability (CIN) increases during replicative aging

Considering the observed depletion of multiple heterochromatin factors in aged yeast cells, as well as the CIN phenotype of a *sir2*Δ mutant (Choy *et al*. 2011), we next tested whether aged cells had a CIN phenotype that was exacerbated by complete loss of *SIR2* and/or *HST1*. Strains utilized for this experiment have an artificial chromosome III bearing a suppressor tRNA gene, *SUP11* (Spencer *et al*. 1990). Loss of the chromosome prevents suppression of an ochre stop codon in *ade2-101*, resulting in the classic *ade2* red colony phenotype. The frequency of nondisjunction events was measured by counting half-sectored red/white colonies from young and aged cell populations (Figure 1G). Sectoring was elevated in young populations of *sir2*Δ and *hst1*Δ mutants, and additively increased in a *sir2Δhst1*Δ double mutant (Figure 1H, black bars, left side of panel). Interestingly, sectoring was significantly higher for aged populations of each strain (Figure 1H, gray bars, left side of panel), suggesting additional age-associated factors were involved. We next tested whether the *sir2*Δ effect on sectoring was related to a pseudodiploid phenotype caused by derepression of the *HM* loci. This reporter strain background was *MAT*α, so we deleted *HMR* (chrIII 293170-294330) to eliminate the a1/a2 transcription factors. Reversal of the pseudodiploid phenotype was confirmed by restoration of mating to the *sir2Δ hmr*Δ strains (data not shown). Importantly, this manipulation significantly suppressed sectoring of the young *sir2*Δ and *sir2Δ hst1*Δ mutants, but not the *hst1*Δ mutant (Figure 1H, middle of panel), indicating there was indeed a *sir2*Δ-induced pseudodiploid effect that contributed to mini-chromosome loss (Figure 1H, black bars, middle of panel). However, aging still increased sectoring in each strain even when *HMR* was deleted (Figure 1H, gray bars, middle of panel), suggesting that the aging-associated CIN factor was unrelated to mating type control.

Because of the observed Net1 depletion (Figure 1E), we hypothesized that age-induced mini-chromosome loss could be related to rDNA instability caused by loss of the RENT complex. To address this idea, *FOB1* was deleted from WT, *sir2*Δ, *hst1*Δ, and *sir2Δ hst1*Δ reporter strains to stabilize the rDNA, followed by retesting the sectoring phenotypes. As shown in Figure 1H (right side of panel), frequency of sectoring observed for each aged *fob1*Δ strain was generally similar to that observed with young *FOB1*^+^ versions of the strains (Figure 1H, black bars, left side of panel), suggesting that destabilization of the rDNA during aging does contribute to the instability of other chromosomes. The *hmr*Δ and *fob1*Δ mutations were potentially suppressing chromosome loss through independent mechanisms, thus begging the question of whether combining them would fully suppress CIN in a *sir2Δ hst1Δ hmrΔ fob1*Δ quadruple mutant. Remarkably, aged cells from this mutant combination lost the mini chromosome marker at a very low rate comparable to young WT cells, with no statistical difference (Figure 1H, 1-way ANOVA).

### Cohesin levels are depleted in aged yeast cells

The above results raised the question of what factor(s) related to rDNA stability, chromosome segregation, and Sir2, was becoming defective in aged cells. Cohesin perfectly fit this profile and was earlier shown in mammalian oocytes to become depleted with age (reviewed in (Jessberger 2012)). To test whether cohesin was another Sir2-linked factor depleted during yeast aging, we C-terminally myc-tagged the Mcd1 or Smc1 subunits (Figure 2A) in the MEP strain background. Western blotting demonstrated that both subunits were significantly depleted from aged cells (Figure 2B), implying depletion of the whole cohesin complex. We also observed depletion of the cohibin/monopolin subunit Lrs4 (Figure S2), which can function as a cohesin clamp at the rDNA (Huang *et al*. 2006). Furthermore, a Myc-tagged Scc2 subunit of the Scc2/Scc4 cohesin loading complex was age-depleted (Figure 2C), though not as severely as the cohesin complex, predicting that the remaining cohesin complex could still be loaded onto chromatin in aged cells. ChIP assays for Mcd1-myc in MEP cells aged for 24 hr demonstrated strong depletion from the rDNA intergenic spacer (IGS1) (Figure 2D), but was surprisingly enriched at the centromeres of chromosomes XI and III. This effect did not appear to significantly extend into the pericentric region of chromosome III (Figure 2E). In order to rule out primer specificity or chromosome size effects, we tested three centromeres from other chromosomes and observed the same trend for each of them (Figure 2F). To confirm the enrichment was centromere specific, we tested two additional sites on chromosome IV that were 15kb and 200kb away from the centromere, and observed no increase in the aged samples compared to young (Figure 2G). To test if the enrichment of Mcd1-myc at centromeres in aged cells was specific to cohesin, and not a ChIP artifact, we next tested the distribution of Sir4-myc, which was also depleted from aged cells (Figure 1C). In this case, Sir4-myc was depleted from *TELXV* in aged cells (one of its normal targets) without any apparent redistribution to centromeres (*CEN4*) or the rDNA (Figure 2H), indicating that not all age-depleted proteins become enriched at centromeres. Based on these results, we hypothesized that as cohesin starts depleting during replicative aging, its association with the rDNA is severely affected while a significant portion of the remaining complex is retained and potentially redistributed to centromeres. This is consistent with an earlier finding that cohesin preferentially associates with pericentromeric regions instead of chromosome arms when Mcd1 expression is artificially reduced below 30% of normal (Heidinger-Pauli *et al*. 2010). We next asked if the centromere enrichment would be lost in an older cell population by extending growth of the MEP culture time course to 36 hr. As shown in Figure 2I, the Mcd1-Myc association with *CEN4* and *CEN14* returned to the original levels observed in young cells, while remaining depleted at the rDNA array (Fig 2J). It is important to note that in a recent report (Pal *et al*. 2018), cohesin enrichment at centromeres was actually reduced in very old yeast cells (>25 generations). Combined with our findings, a model emerges whereby cohesin enrichment at centromeres is initially enhanced during aging, but then catastrophically lost as cells approach the end of their lifespan.

**Figure 2.**
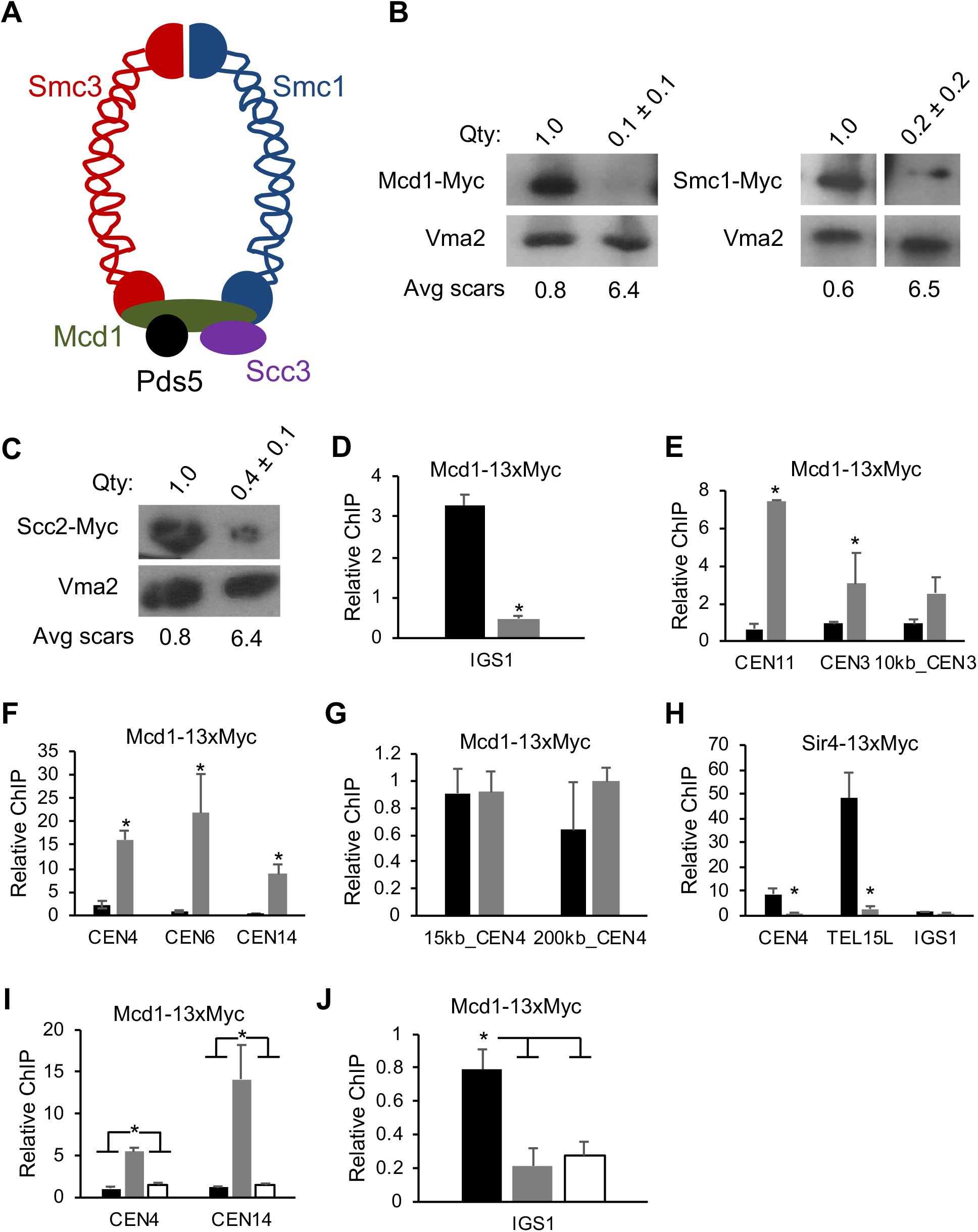
Chromosome instability is linked to rDNA stability and cohesin redistribution. **(A)** Schematic of the cohesin complex subunits. **(B)** Western blot of Mcd1-13xMyc and Smc1-13xMyc from young and aged cells. **(C)** Western blot of 13xMyc tagged Scc2. **(D, E, F, G)** ChIP-qPCR of Mcd1-13xMyc in young cells (black bars) and aged cells (gray bars) at the indicated loci normalized to background signal in the *STE2* ORF. **(H)** ChIP-qPCR of Sir4-13xMyc in young and aged cells normalized to an intergenic *PDC1* site that shows SIR complex association (Li *et al*. 2013). **(I, J)** ChIP-qPCR of Mcd1-13xMyc from a time course of young, 24 hr, and 36 hr aged cells at indicated loci normalized to *STE2* background signal. Asterisks indicate significant differences between young and aged cells (p<0.05).

### Sister chromatid cohesion is compromised in aged yeast cells

Previous work found that SCC was surprisingly normal despite the forced reduction (<30%) of Mcd1 protein levels (Heidinger-Pauli *et al*. 2010), leading us to ask whether cohesion would be maintained in our moderately aged yeast cells that were also depleted for cohesin, yet still showed enrichment at centromeres. To this end, we utilized strains with a LacO array located approximately 10 kb away from centromere IV (*CEN4*) as a proxy for centromeric cohesion, or at the *LYS4* locus on chromosome IV, located approximately 400 kb away from the centromere, as a proxy for arm cohesion (Unal *et al*. 2008; Guacci and Koshland 2012). Differential positioning of the array had no significant impact on RLS (Figure S3). SCC was monitored by LacI-GFP appearing either as one dot in the case of cohesion maintenance or two dots in the case of cohesion loss. Using an *mcd1-1* temperature sensitive mutant as a positive control (Guacci *et al*. 1997), we observed a large increase in two dots when cells were synchronized in mitosis with nocodazole and shifted to 37°C (Figure 3A and B). WT cells for the equivalent reporter strain were next biotinylated and aged for 24 hours, followed by purification with magnetic streptavidin beads and arrest with nocodazole. Analyzing cells with >7 bud scars, which is older than the average bud scar count of our western blot experiments, revealed a mild loss of cohesion (Figure 3C and D). A similar analysis was then performed without the use of nocodazole to rule out any side-effects related to triggering the mitotic spindle checkpoint. To avoid misinterpreting anaphase events as lost SCC, we C-terminally tagged the spindle pole body subunit Spc42 with dsRED, and only counted GFP dots from large-budded cells where the spindle pole bodies had not separated between the mother and daughter cell. With this analysis, the fold-change of SCC defect between young and aged cells was more pronounced (Figure 3E and F), but the absolute frequency of loss was still much weaker than observed with the *mcd1-1* mutant, which was not surprising given that mean lifespan of this strain background is ∼16 generations (Figure S3). In very old cells (>25 bud scars) of a different strain background, an independent study observed SCC loss on chromosome XII at a frequency that approached 50% (Pal *et al*. 2018) suggesting that CIN may become most severe in the oldest cells where cohesin is not only depleted from the rDNA, but also from centromeres.

**Figure 3.**
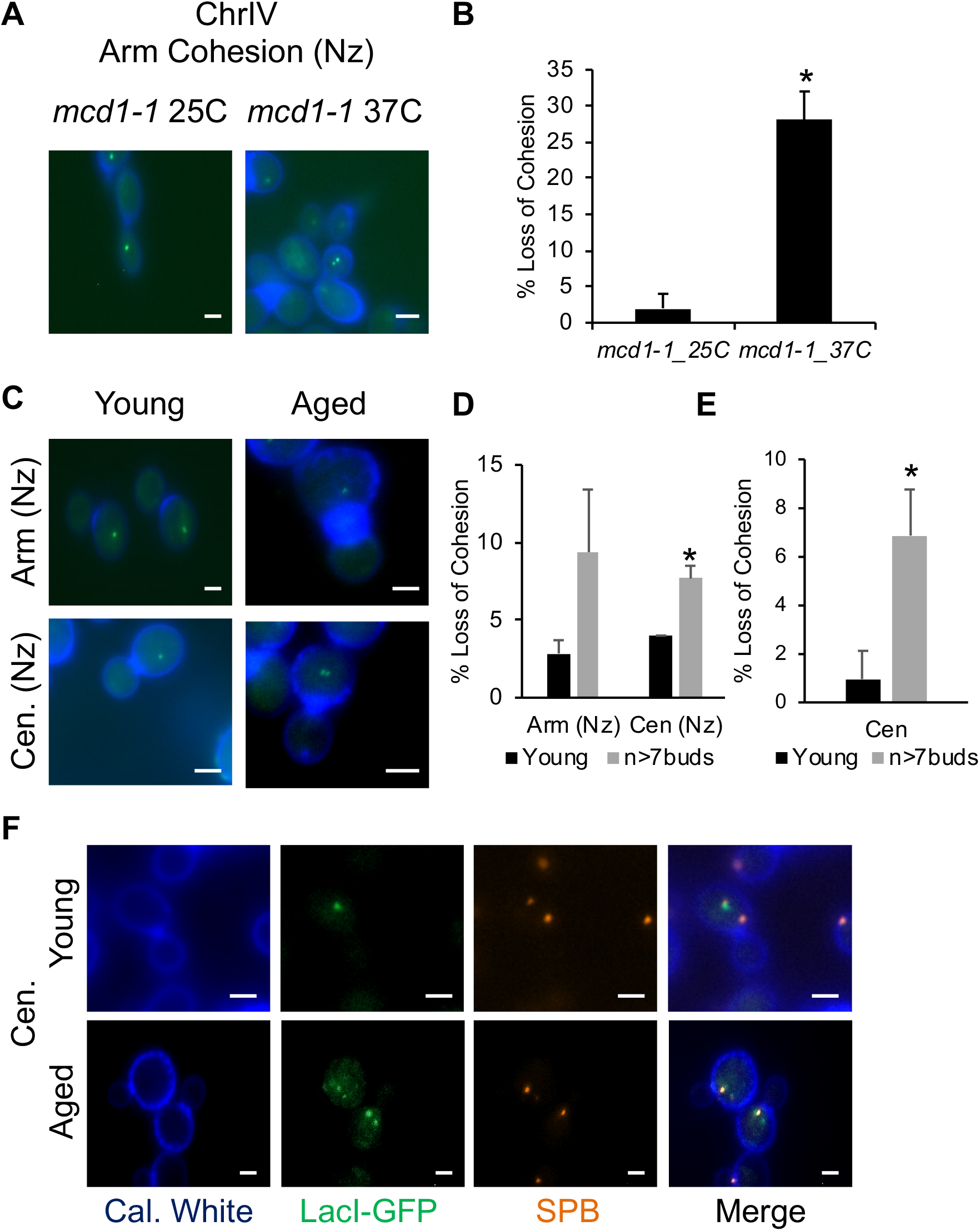
Sister chromatid cohesion is weakened in aged yeast cells. **(A)** Representative control images of arm cohesion in an *mcd1-1* mutant at permissive (25°C) and non-permissive (37°C) temperatures. **(B)** Quantification of cohesion maintenance (1-dot) or loss (2-dots) from 100 *mcd1-1* cells. **(C)** Representative images of young (log-phase) or aged cells arrested with nocodazole (Nz) and monitored for centromere or arm cohesion. **(D)** Quantifying cohesion loss (2-dots) in cells with at least 7 bud scars. *p<0.05, two-tailed t-test (n>50 cells). Quantification of cohesion loss for young and aged cells in the absence of nocodazole **(F)** Representative images of nocodazole untreated cells. White scale bar represents 2 microns.

### RLS is modulated by Mcd1 expression levels

Since Sir2 and cohesin are both naturally depleted from replicatively aging yeast cells (Figures 1A and 2B), and mild Sir2 overexpression extends RLS (Kaeberlein *et al*. 1999), we hypothesized that manipulating cohesin expression levels would also impact CIN and RLS in a dose-dependent manner. We initially attempted to overexpress the Mcd1 subunit from a galactose inducible *GAL1* promoter and then measure mini-chromosome loss frequency by counting ½ sectored colonies. However, simply growing the reporter strain in galactose-containing media, even with an empty expression cassette, resulted in severe mini-chromosome loss compared to glucose-containing media (Figure 4A and B). This effect was specific to galactose, as growth with another non-preferred carbon source (raffinose) had no effect on sectoring (Figure 4A and B). Though not useful for assaying the effects of *MCD1* overexpression, it was still possible that the unexpectedly high CIN phenotype would correlate with reduced RLS. We therefore measured RLS with the mini-chromosome reporter strain on YEP plates with 2% glucose, galactose, or raffinose. As shown in Figure 4C, galactose strongly decreased the mean RLS by ∼50% compared to glucose (9.2 vs. 18.9 divisions, p<1.0×10^−15^), while raffinose only had a marginal effect (15.5 divisions, p<0.01). To confirm the galactose effect on RLS was not specific to the mini-chromosome strain, lifespan assays were repeated with the commonly used strains BY4741 (*MAT*a) and BY4742 (*MAT*α). Again, a significant decrease in mean lifespan was observed for BY4741 (17.7 divisions) and BY4742 (18.9 divisions) on galactose as compared to glucose (24.3 and 24.2 divisions, respectively) (Figure S4, p<0.001), suggesting that galactose triggers a high rate of CIN through an unknown mechanism that also shortens RLS.

**Figure 4.**
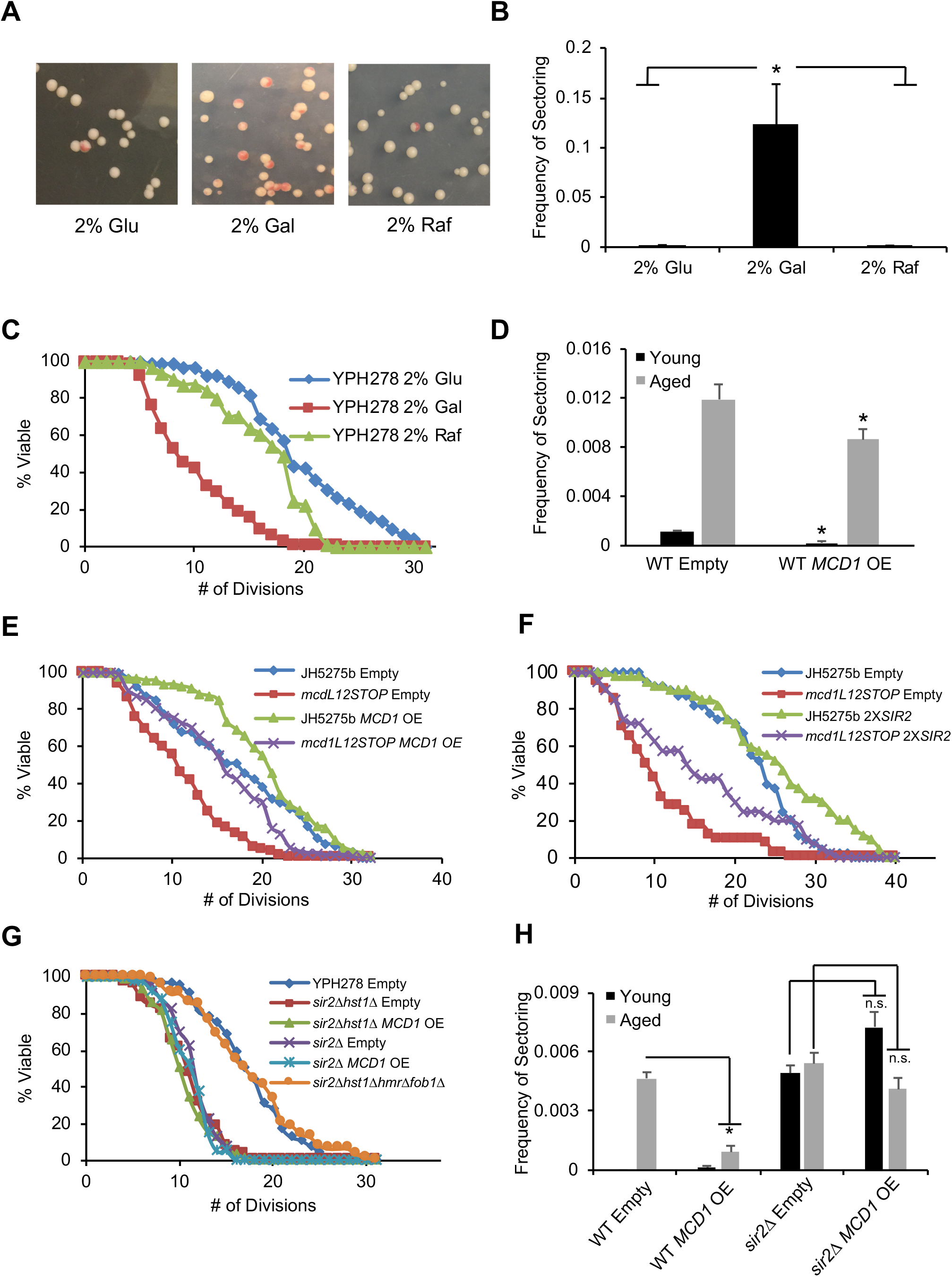
Modulation of RLS and CIN by manipulating *MCD1* and *SIR2* expression levels. **(A)** Representative images of chromosome loss (sectoring) for WT (YPH278) cells grown continuously in 2% glucose, galactose, or raffinose. **(B)** Quantification of half-sector colonies shown in 4A. *p<0.001, two-tailed student t-test. **(C)** RLS assay of YPH278 cells growing on rich YEP agar plates containing either 2% glucose, galactose, or raffinose. (n=64, each condition; mean RLS × ♦18.9, ■ 9.2, ▲ 15.5) **(D)** Quantification of chromosome loss (1/2 sectors) for strains with an integrated tet^On^ empty (pRF10) or *MCD1* (pRF11) construct. RLS assays of WT (*MCD1*^+^) and *mcd1L12STOP* strains with integrated pRF10 or pRF11 constructs (mean RLS × ♦16.2, ■ 10.3, ▲ 19.2, ×15.6). **(F)** RLS assay of WT and *mcd1L12STOP* strains with integrated empty, pRS306, or *SIR2* containing plasmid, pJSB186 (n=40 cells each; mean RLS × ♦21.5 ■ 9.9 ▲ 30.4 ×14.8). **(G)** RLS assay with *MCD1* OE or *fob1Δ hmr*Δ rescue of *sir2*Δ or *sir2Δ hst1*Δ mutants (mean RLS × ♦16.5, ■ 10.1, ▲ 9.7, ×10.7, ∗ 10.25, • 16.7). **(H)** rDNA recombination (marker loss, ½ sector) assay with rDNA marker loss reporter strain W303AR bearing *ADE2* within the rDNA array. (* p<0.05, two-tailed t-test).

To circumvent the use of galactose for *MCD1* overexpression we turned to an inducible “Tet-On” promoter that is activated by doxycycline (Belli *et al*. 1998). Strains harboring this integrated cassette transcriptionally overexpressed *MCD1* approximately 2- to 7-fold compared to the empty vector control (Figure S5A and B). In the mini-chromosome reporter strain, *MCD1* overexpression significantly reduced the frequency of ½ sectored colonies in both young and aged populations (Figure 4D), in agreement with *MCD1* being isolated as a high copy suppressor of CIN using a different reporter system (Zhu *et al*. 2015). Next, *MCD1* was overexpressed in a strain containing an ochre stop codon in the *MCD1* open-reading frame (*mcd1L12STOP*) that reduces Mcd1 protein to ∼30% of normal (Heidinger-Pauli *et al*. 2010). With an empty pCM252 control *CEN* vector, *mcd1L12STOP* exhibited a 40% reduction in mean RLS (9.9 divisions) compared to an isogenic WT control (16.5 divisions), indicating that Mcd1 depletion shortens RLS (Figure 4E, p<0.001). Overexpressing *MCD1* almost fully restored longevity to the mutant (14.7 divisions) and, remarkably, also extended RLS of the control WT strain to 19.6 divisions (p<1.0×10^−7^), which was primarily due to improved survival during the first ∼15 divisions and then followed by a steeper decline in viability (Figure 4E). This biphasic pattern was highly reproducible and suggested that the *CEN MCD1* OE plasmid was being lost around mid-life due to increased chromosome loss, as previously seen in aged cells of our CIN assay (Figure 1H). To account for this potential variable, we also integrated the empty and *MCD1* OE vectors into a different FY834 strain background related to the long lived BY4741/42 background (Winston *et al*. 1995; Brachmann *et al*. 1998). Not only did *MCD1* OE extend the mean RLS (28.0 divisions) in this background compared to the control (21.0 divisions), but the maximum number of divisions was also increased by 25% (Figure S6, p<1.0×10^−7^). We therefore conclude that similar to Sir2, Mcd1 is a strong dose-dependent longevity factor.

Sir2 functions upstream of cohesin for RLS (Kobayashi *et al*. 2004; Kobayashi and Ganley 2005), and for SCC at *HMR* (Wu *et al*. 2011), implying that Sir2 is upstream of cohesin loading and function. However, this relationship could potentially be more complex since both factors are depleted with age. To explore this further, we next tested if *SIR2* overexpression could rescue the short RLS of an Mcd1-depleted *mcd1L12STOP* strain by integrating a second copy of *SIR2* (2x*SIR2*) at the *LEU2* locus. As shown in Figure 4F, 2x*SIR2* partially rescued mean RLS of the *mcd1L12STOP* strain (14.1 versus 9.9 divisions, p<1.0×10^^−4^), and also increased maximum RLS of the WT strain. Reciprocally, we asked if *MCD1* overexpression would suppress the short RLS of a *sir2*Δ or *sir2Δ hst1*Δ mutant. The double mutant was included to rule out any redundancy between the two sirtuins. Mean RLS was clearly not increased by *MCD1* overexpression as compared to empty vector for either the single (10.7 versus 10.2) or double mutant (10.1 versus 9.7 divisions, Figure 4G), confirming that *SIR2* was required for *MCD1* in regulating RLS. Interestingly, the *sir2Δ hst1Δ fob1Δ hmr*Δ quadruple mutant combination, which suppressed CIN in aged cells (Figure 1H), fully restored RLS of the short lived *sir2Δ hst1*Δ combination to WT levels (Figure 4G, p=n.s.), suggesting that CIN is a key driver of replicative aging downstream of Sir2 and Hst1.

Considering the strong depletion of cohesin from rDNA in aged cells (Figure 2D), and extended RLS when *MCD1* was overexpressed (Figure 4E), we next tested if age-induced rDNA instability was suppressed by *MCD1* overexpression using a reporter strain harboring *ADE2* in the rDNA array (Kaeberlein *et al*. 1999). There was a large increase of red/white ½ sectoring (marker loss) from aged cells that was suppressed upon *MCD1* overexpression (Figure 4H). In the absence of *SIR2*, ½ sectoring was high from young and aged cells when the empty vector (pRF10) was integrated, and *MCD1* overexpression did not significantly reduce rDNA instability in either population (Figure 4H), indicating that at least some Sir2 was required for Mcd1 to impact rDNA stability. We conclude that loss of Sir2 and cohesin in aging cells causes rDNA array instability that generally exacerbates CIN.

### RLS extension by CR correlates with improved chromosome stability

Reducing glucose concentration in the growth medium is effective at extending RLS and is considered a form of caloric restriction (CR) for yeast (Jiang *et al*. 2000; Lin *et al*. 2002). There have been several hypotheses put forth for the underlying mechanisms, including stabilization of the rDNA, as CR suppresses recombination within the rDNA (Lamming *et al*. 2005; Riesen and Morgan 2009; Smith *et al*. 2009). Since *MCD1* overexpression suppressed rDNA recombination and extended RLS, we hypothesized that CR may suppress the abbreviated RLS of a cohesin-depleted *mcd1L12STOP* mutant strain. Indeed, CR extended RLS of both the WT and *mcd1L12STOP* strains (Figure 5A, p<0.01 and p<1.0×10^−7^ respectively). However, the suppression was apparently not due to maintenance of global cohesin levels because steady state Mcd1-13xMyc was still depleted in glucose restricted aged cells (Figure 5B). CR also strongly suppressed minichromosome loss in young and aged cells, even in the *sir2Δ hst1*Δ double mutant (Figure 5C). Importantly, this CR effect also correlated with almost complete rescue of RLS for the *sir2Δ hst1*Δ mutant (Figure 5D, p=n.s.). Taken together, the results support a mechanism for RLS extension by CR, whereby stabilization of the rDNA locus helps maintain general mitotic chromosome stability to protect against aneuploidy.

**Figure 5.**
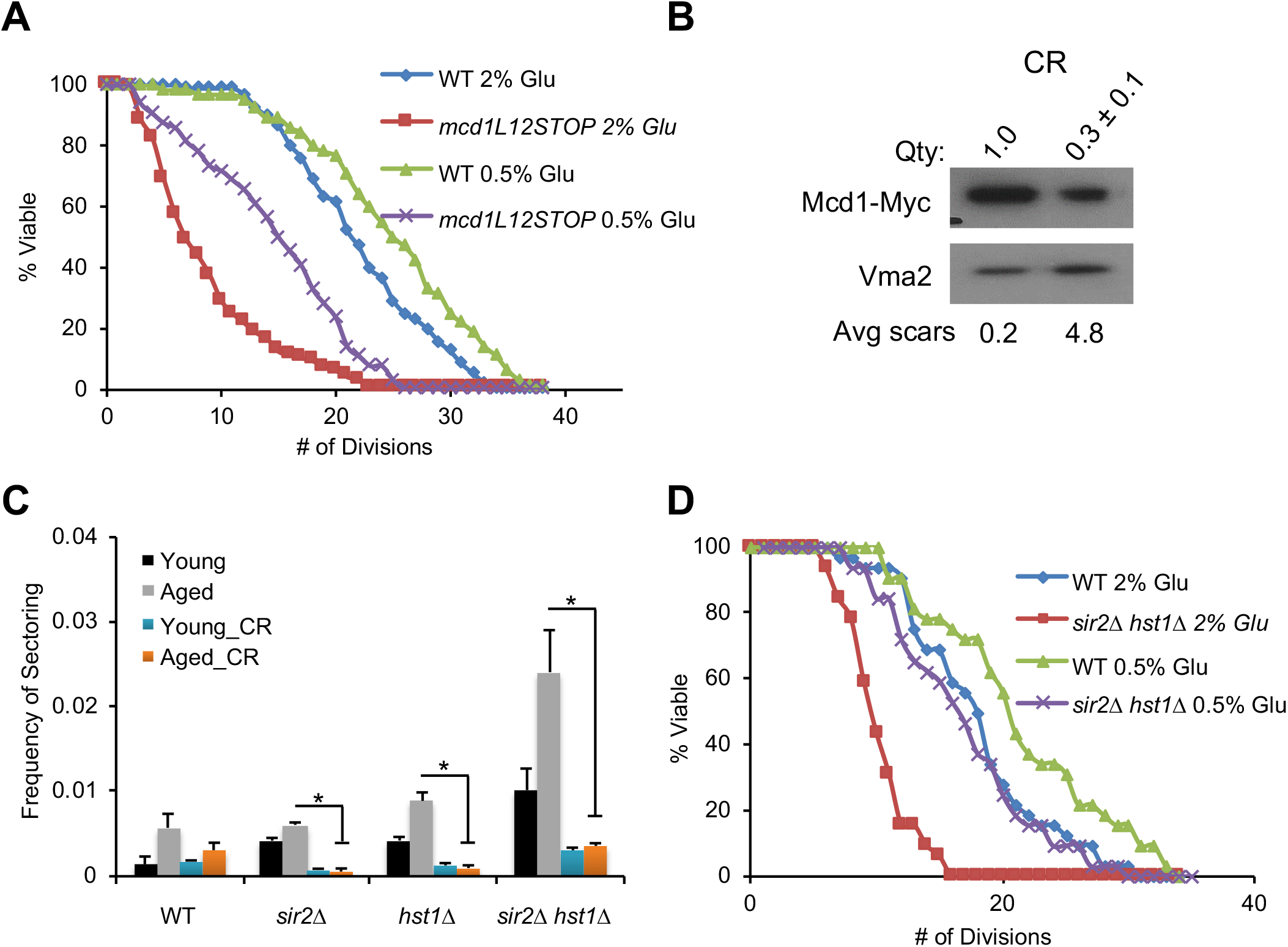
CR suppresses the CIN and RLS defects of *sir2*Δ and *mcd1* mutants. **(A)** RLS assay of WT (JH5275b) and *mcd1L12STOP* (JH5276b) strains under normal 2% glucose and CR (0.5% glucose) conditions (n=32 cells; mean RLS × ♦21.3, ■ 8.1, ▲ 24.1, ×13.8). **(B)** Western blot of Mcd1-13xMyc for cultures grown in YEP media containing 0.5% glucose. **(C)** Chromosome loss (sectoring) assay of strains grown in media containing 0.5% glucose. Data from Figure 1H included for reference. *p<0.05, two-tailed t-test. **(D)** RLS assay of WT and *sir2Δhst1*Δ mutant strains under normal and CR conditions. (n=32 cells; mean rls × ♦19.6, ■ 9.3, ▲ 23.5, ×17.1).

### The rDNA array has an opportunity to interact with centromeres during anaphase

To conclude this study, we asked whether there is any mechanistic connection between the rDNA and centromeres that could cause CIN. If rDNA instability has a direct effect on SCC during aging, then we should observe elevated dissociation of sister chromatids in a *sir2*Δ mutant and improved SCC in a *fob1*Δ mutant. However, as shown in Figure S7, the frequency of 2 GFP dots in the SCC assay for these two mutants in aged populations was comparable to WT (see Figure 3E and F). Alternatively, the rDNA could potentially affect centromere function through direct contacts. Previous Hi-C analysis of the yeast genome and fluorescence microscopy of nucleolar proteins positioned the rDNA off to one side of the nucleus, apparently secluded from the rest of the genome (Gotta *et al*. 1997; Duan *et al*. 2010). The repetitive nature of rDNA precludes it from appearing in Hi-C contact maps, but closer inspection of iteratively corrected chromosome XII contact maps at 10 kb resolution indicated a clear interaction between unique sequences flanking the centromere-proximal (left) edge of the array and *CEN12* (Figure 6A, yellow arrow). We hypothesized that this contact was regulated by Sir2 since it is in the vicinity of a known tRNA boundary (tQ(UUG)L) for rDNA silencing (Biswas *et al*. 2009), but deletion of *SIR2* had no effect on the contact (data not shown). Interestingly, all centromeres of the yeast genome, including *CEN12*, cluster together in asynchronous cell population Hi-C data (Figure 6B, yellow arrows; (Duan *et al*. 2010)), which potentially places them in proximity with the rDNA given the association of *CEN12* with sequences flanking the rDNA. During anaphase, the rDNA is thought to be separated from centromeres, but analysis of Hi-C data for cell-cycle synchronized *cdc15-2*, which arrests cells in anaphase, revealed that the rDNA-proximal/*CEN12* contact specifically occurs during anaphase (Figure 6C, (Lazar-Stefanita *et al*. 2017)), during which time the centromeres are still clustered together by the spindle pole body (Figure 6D).

**Figure 6.**
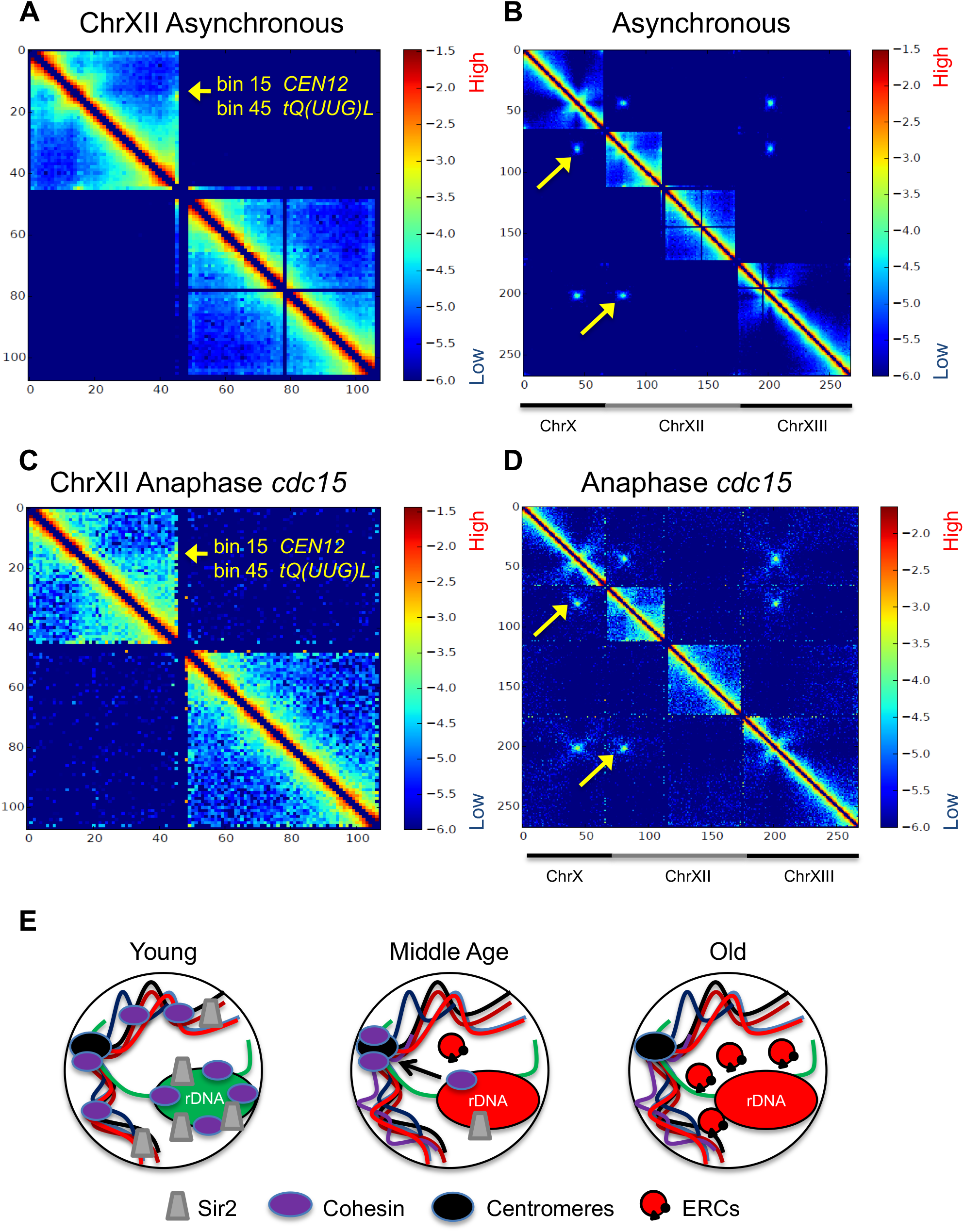
Chromosomal instability during replicative aging is linked to the rDNA. **(A)** Iteratively corrected and read-normalized heatmap of ChrXII Hi-C contact data at 10 kb resolution in WT cells growing asynchronously, revealing an interaction between *CEN12* (bin 15) and unique sequence adjacent to the rDNA (bin 45), indicated by yellow arrow. **(B)** Contact heatmap of chromosomes XI, XII, and XIII showing centromere clustering in asynchronously growing WT cells. Yellow arrows indicate examples of centromere alignment with *CEN12* and sequence adjacent to the rDNA. **(C)** Iteratively corrected Chromosome XII heatmap from *cdc15* cells arrested in anaphase. Yellow arrow indicates the interaction between CEN12 and the rDNA-adjacent bin. **(D)** Contact map showing interactions between centromeres in the anaphase-arrested *cdc15* mutant. **(E)** Model for cohesin and Sir2 depletion from the rDNA of aged mother cells. Cohesin is initially retained at centromeres (Middle Age, black arrow), and then ultimately lost in the oldest cells. ERCs begin to form during middle age and may contribute in a self-perpetuating cycle resulting in chronic destabilization of the rDNA array due to loss of cohesin, RENT, and the cohibin complex (red).

Taken together, these results suggest that the rDNA may transiently contact the centromeres during mitosis, providing a potential window of time for a destabilized rDNA array to negatively impact the integrity of general chromosome segregation during mitosis (Figure 6E).

## Discussion

### Nuclear protein depletion during replicative aging as a paradigm for aging pathologies

During this study the majority of chromatin-associated proteins that we analyzed by western blotting were depleted in replicatively aged yeast cells. The only protein unaffected by age that we tested, other than the vacuolar Vma2 control, was Sir3 (Figure 1D). A similar proportion of homologous recombination proteins were depleted in an independent analysis of aged cells, with Rad52 the only one tested that was not affected (Pal *et al*. 2018). These results suggest there is at least some selectivity to the depletion of nuclear proteins in aged cells. However, the large number of depleted factors also makes it likely that targeted nuclear protein deficiency could lead to multiple age-associated phenotypes. Replicatively aging yeast cells appear especially susceptible to this phenomenon, as even total core histone levels are depleted (Hu *et al*. 2014). Evidence also exists for histone depletion during aging of metazoan organisms, including mammals (reviewed in (Song and Johnson 2018)). More generally, global protein turnover is elevated in cells from prematurely aging progeria patients, which may trigger higher translation rates (Buchwalter and Hetzer 2017). This is significant because reducing translation is a means of extending lifespan in multiple organisms (Mehta *et al*. 2010). The mechanism(s) driving nuclear protein depletion in aged yeast mother cells or other organisms remain unclear.

The specificity for Sir2/Sir4 depletion over Sir3 is intriguing given that Sir2 and Sir4 form a tight complex that allosterically stimulates the deacetylase activity of Sir2 (Hsu *et al*. 2013), while Sir3 is a subunit of the SIR holocomplex (Oppikofer *et al*. 2011). Since Sir3 levels are elevated in aged cells (Figure 1D; (Janssens *et al*. 2015)), it is likely to have a function independent of the SIR complex during aging. The mechanism for Sir2/Sir4 depletion from aged cells remains uncharacterized, though in non-aging cell populations the stability/turnover of Sir4, but not Sir2, is mediated by the E3 ubiquitin ligase San1 (Dasgupta *et al*. 2004), which has also been implicated as a quality control E3 ligase for mutated/unfolded nuclear proteins (Gardner *et al*. 2005). Whether San1 controls Sir4 stability during aging remains unknown, but since Sir4 is more severely depleted than Sir2 in aged cells (Figures 1A and 2C), Sir4 could be selectively depleted from the SIR complex, thus leaving Sir2 unprotected and subject to turnover through a different mechanism. Consistent with this idea, we note that Sir4 is depleted independently of Sir2 during extended G1 arrest (Larin *et al*. 2015). Alternatively, Sir2/Sir4 could be equally depleted as a complex from telomeres and the *HM* loci (not necessarily via San1), leaving the nucleolar pool of Sir2/RENT as more resistant to aging. Under this scenario, protecting integrity of the rDNA array could take precedence over other heterochromatic domains. Interestingly, the *Schizosaccharomyces pombe* San1 ortholog has also been implicated in a chaperone-assisted degradation pathway that functions in quality control of kinetochores to promote chromosome stability (Kriegenburg *et al*. 2014).

Sir2 depletion in replicatively aged yeast cells is reminiscent of Sirt1 depletion in serially passaged mouse embryonic fibroblasts (MEFs), which correlates with declining mitotic activity (Sasaki *et al*. 2006). Sir2 and Sirt1 are both known to function in regulating DNA replication origins (Hoggard *et al*. 2018; Utani and Aladjem 2018), and the effect of deleting *SIR2* on early origin firing is thought to be mediated by competition for limiting factors with the repeated rDNA origins (Yoshida *et al*. 2014). Furthermore, CR has been proposed to extend RLS by reducing rDNA origin firing, which improves overall genome replication (Kwan *et al*. 2013). This may help explain why CR can extend RLS and suppress CIN even when *SIR2* and *HST1* are deleted. Conversely, depleted cohesin in old cells could potentially cause rDNA instability by impacting DNA replication and repair.

### A precarious balance between rDNA and centromeric cohesin

SCC ensures chromosomes are not segregated until the Mcd1/Scc1 cohesin subunit is cleaved in response to a mitotic spindle checkpoint signal that all chromosomes are properly attached to microtubules and aligned at the metaphase plate (reviewed in (Marston 2014)). Cohesin is also critical during meiosis, and it is well established in mammals that SCC defects occur in the oocytes of older mothers, causing meiotic chromosome missegregation events during both anaphase I and II (Jessberger 2012). This phenomenon is believed to be a major mechanism for increased aneuploidy risk that usually results in embryonic lethality, or in the case of chromosome 21 trisomy, Down’s syndrome. The meiotic cohesin subunit Rec8 is depleted in the oocytes of older mice, as is Shugoshin (Sgo2), which normally protects/maintains centromeric cohesin (Lister *et al*. 2010). More recent experiments in *Drosophila* suggest that oxidative stress in aged oocytes contributes to the SCC defects (Perkins *et al*. 2016). Our results in replicatively aging yeast cells reveal that aging-induced cohesin depletion and the resulting chromosome missegregation can extend to mitotic cells. Though cohesin depletion or defects have not been reported for aging mammalian somatic cells, the mitotic spindle checkpoint protein BubR1 is depleted in dynamic somatic tissues such as spleen in aged mice (Baker *et al*. 2004). Deficiency of this protein results in premature aging phenotypes (Baker *et al*. 2004), while overexpression extends lifespan (Baker *et al*. 2013). This is similar to the effects we observe with Mcd1 depletion and overexpression on yeast RLS. Interestingly, BubR1 is also a deacetylation target of Sirt2, which appears to stabilize the protein and extend lifespan, thus linking mitotic spindle checkpoint regulation to NAD^+^ metabolism (North *et al*. 2014). It remains unclear if Sir2, Hst1, or other sirtuins regulate the yeast BubR1 ortholog, Mad3, or additional checkpoint and kinetochore proteins.

SCC is the canonical function for cohesin, though the complex also functions in establishing and regulating genome organization at the level of chromatin structure, gene regulation, and double strand break (DSB) repair (reviewed in (Uhlmann 2016)). Among these various processes, SCC at centromeres appears the most critical because artificial depletion of Mcd1 to <30% of normal levels results in preferential cohesin binding to pericentromeric regions rather than cohesin associated regions (CARs) on chromosome arms (Heidinger-Pauli *et al*. 2010). SCC was also well maintained in these strains at the expense of normal chromosome condensation, DNA repair, and rDNA stability (Heidinger-Pauli *et al*. 2010). In aged yeast cells, we observed relative enrichment of Mcd1-myc at centromeres as compared to loss at the rDNA (IGS1) locus (Figures 2D and 2E) consistent with pericentromeric cohesin retention in the artificially depleted system. Despite maintaining the cohesin complex at centromeres, SCC was still slightly impaired in the aged cells, but only if we analyzed cells >7 generations old. We suspect cohesin was reduced at centromeres in these older cells, which would be consistent with the loss of centromeric Mcd1 enrichment when cells were aged longer for 36 hr, but it was also likely that some of the centromere-localized Mcd1 was non-functional. These results are in line with an independent study that analyzed significantly older mother cells (∼25 generations) and observed reduced cohesin enrichment at centromeres and severe loss of SCC (Pal *et al*. 2018). Another recent study using single cell microfluidics found that chromosome loss was very common just prior to the last cell division (Neurohr *et al*. 2018). Collectively, the results suggest that centromere-associated cohesin is preferentially retained during the initial stages of replicative aging, but then eventually breaks down below a critical threshold in the oldest cells.

Numerous nuclear proteins are depleted in aged yeast cells, not just cohesin subunits, so we hypothesize that defects in other nuclear processes mediated by such factors also contribute to SCC defects and chromosome instability either directly or indirectly. The depleted cohesin loading complex (Scc2/4) is an obvious candidate due to its role in loading cohesin at centromeres and CARs. Similarly, the depleted cohibin complex (Figure S2) is proposed to act as a cohesin clamp onto rDNA chromatin (Huang *et al*. 2006), and also functions at centromeres to maintain mitotic integrity (Bitto *et al*. 2015). Sir2 and Hst1 are also obvious candidates given the earlier finding that H4K16 deacetylation at centromeres by Sir2 helps maintain chromosome stability (Choy *et al*. 2011). Part of this effect is apparently due to the pseudo-diploid phenotype of a *sir2*Δ mutation, which has been previously shown to impact RLS (Kaeberlein *et al*. 1999). Hst1 also binds centromeric DNA *in vitro* and *in vivo* (Ohkuni and Kitagawa 2011), though the functional relevance of that association remains uncharacterized. The suppression of age-associated mini-chromosome loss in the absence of *FOB1* clearly points to rDNA instability as an unexpected source of general CIN. Such a relationship is reinforced by the observed depletion of nucleolar proteins Net1 and Lrs4 in aged cells (Figures 2E and S2), both of which are required for normal rDNA/nucleolar integrity and stable cohesin association with the rDNA (Smith *et al*. 1999; Straight *et al*. 1999; Huang *et al*. 2006).

How could destabilization of the rDNA locus result in general chromosome instability and shortened RLS? As depicted in Figure 6, unique sequence flanking the rDNA contacts the centromere of chromosome XII, thus placing it in proximity to other centromeres during anaphase. Whether the actual rDNA genes contact centromeres remains unclear due to the current limitations of Hi-C analysis with repetitive DNA. However, specific regions of the rDNA were previously shown to associate with various non-rDNA chromosomal regions using an anchored 4C approach (O’Sullivan *et al*. 2009). Furthermore, multiple nucleolar associated domains have been identified in metazoan cells that copurify with nucleoli (Matheson and Kaufman 2016). Loss of cohesin from the rDNA could potentially disrupt long-range interactions with centromeres or non-centromeric regions of cohesin association that influence chromosome integrity. One potential mechanism could be significant disruption of overall chromosome condensation during mitosis, as cohesin appears to play a larger role in the DNA looping associated with chromosome condensation in budding yeast than previously thought (Schalbetter *et al*. 2017).

Interestingly, another class of nuclear factors depleted in aged yeast cells are several DNA repair proteins (Pal *et al*. 2018). Consequently, the lack of proper DNA repair while the rDNA becomes destabilized correlates with fragmentation of chromosome XII and the other chromosomes (Pal *et al*. 2018). Rad52 foci also appear in aged cells indicating persistent DNA damage (Neurohr *et al*. 2018). It was proposed that accumulation of breaks and rearrangements ultimately causes cell death during replicative aging. Such cells were significantly older (>25 divisions) compared to the cells in our study, which exhibited a maximum of 13 divisions after 24 hr. Alternatively, it is possible that these presumably random rearrangements disrupt normal SCC, leading to CIN.

### Aneuploidy as an aging mechanism

All 16 *S. cerevisiae* chromosomes harbor essential genes, so if a single chromosome is lost from a haploid yeast cell, then the affected mother or daughter cell should become inviable and no longer divide. Given the elevated frequency of chromosome loss during replicative aging, the chances of generating an inviable mother cell during a replicative aging assay increases after each subsequent division. Therefore, at least a portion of the replicative lifespan in haploid yeast cells is controlled by the ability to maintain all 16 chromosomes. Complete loss of a chromosome would not be an immediate viability issue for diploid cells, however, due to the chances of losing both homologs in a single mitosis being exceedingly rare. On the other hand, haploid strains that are disomic for individual chromosomes are often short lived, with longer chromosomes typically having larger effects (Sunshine *et al*. 2016). It was hypothesized that such strains suffer from proteotoxic stress due to inappropriate protein expression levels. Therefore, a similar mechanism could shorten RLS in a diploid strain that is trisomic for an individual chromosome, though this has not yet been tested. Aneuploidy is also a hallmark of aging in the germline (Nagaoka *et al*. 2012), and somatic tissues of mammals (Lushnikova *et al*. 2011; Baker *et al*. 2013), making it a conserved feature of aging from yeast to humans.

Another exciting feature of this study is the suppression of CIN by CR growth conditions that extend RLS. This effect was independent of the reduced cohesin levels in aged cells, and even improved RLS of the cohesin-depleted strains. Since SCC is normal in the cohesin-depleted strain (Heidinger-Pauli *et al*. 2010), we hypothesize that CR reinforces other processes that are defective due to reduced cohesin or other depleted factors that promote rDNA stability. Indeed, CR is known to suppress rDNA instability in yeast cells (Riesen and Morgan 2009; Smith *et al*. 2009), and improve overall genome replication efficiency (Kwan *et al*. 2013). Hi-C analysis also suggests there could be direct effects of rDNA structure on centromere function, which will be a focus of future investigation.

## Acknowledgments

We thank Dan Gottschling and all lab members for kindly providing yeast strains and initial advice on the MEP system. Stefan Bekiranov, Job Dekker, Jon Belton, Maitreya Dunham, Ivan Liachko, Maxim Imakev, and Anton Goloborodko all provided valuable advice on Hi-C protocols and analysis methods. We thank Doug Koshland for providing the Mcd1 reduction and cohesion assay strains, and Matt Kaeberlein for rDNA marker loss strains. We also thank Jef Boeke, Marc Gartenberg, and Dan Gottschling for providing antibodies. Special thanks to Todd Stukenberg for microscopy assistance. Lastly, thanks to David Auble for critically reading the manuscript and providing comments prior to submission. We declare no conflicts of interest.

## References

Ausubel, F. M., R. Brent, R. E. Kingston, D. D. Moore, J. G. Seidman et al. (Editors), 2000 Current Protocols in Molecular Biology. John Wiley & Sons, Inc., New York.

Baker, D. J., M. M. Dawlaty, T. Wijshake, K. B. Jeganathan, L. Malureanu et al., 2013 Increased expression of BubR1 protects against aneuploidy and cancer and extends healthy lifespan. Nat Cell Biol 15: 96–102.

Baker, D. J., K. B. Jeganathan, J. D. Cameron, M. Thompson, S. Juneja et al., 2004 BubR1 insufficiency causes early onset of aging-associated phenotypes and infertility in mice. Nat Genet 36: 744–749.

Belli, G., E. Gari, L. Piedrafita, M. Aldea and E. Herrero, 1998 An activator/repressor dual system allows tight tetracycline-regulated gene expression in budding yeast. Nucleic Acids Res 26: 942–947.

Belton, J. M., and J. Dekker, 2015 Measuring chromatin structure in budding yeast. Cold Spring Harb Protoc 2015: 614–618.

Biswas, M., N. Maqani, R. Rai, S. P. Kumaran, K. R. Iyer et al., 2009 Limiting the extent of the *RDN1* heterochromatin domain by a silencing barrier and Sir2 protein levels in *Saccharomyces cerevisiae*. Mol Cell Biol 29: 2889–2898.

Bitto, A., A. M. Wang, C. F. Bennett and M. Kaeberlein, 2015 Biochemical genetic pathways that modulate aging in multiple species. Cold Spring Harb Perspect Med 5.

Brachmann, C. B., A. Davies, G. J. Cost, E. Caputo, J. Li et al., 1998 Designer deletion strains derived from *Saccharomyces cerevisiae* S288C: a useful set of strains and plasmids for PCR-mediated gene disruption and other applications. Yeast 14: 115–132.

Brachmann, C. B., J. M. Sherman, S. E. Devine, E. E. Cameron, L. Pillus et al., 1995 The *SIR2* gene family, conserved from bacteria to humans, functions in silencing, cell cycle progression, and chromosome stability. Genes Dev 9: 2888–2902.

Bryk, M., M. Banerjee, M. Murphy, K. E. Knudsen, D. J. Garfinkel et al., 1997 Transcriptional silencing of Ty1 elements in the *RDN1* locus of yeast. Genes Dev 11: 255–269.

Buchwalter, A., and M. W. Hetzer, 2017 Nucleolar expansion and elevated protein translation in premature aging. Nat Commun 8: 328.

Buck, S. W., C. M. Gallo and J. S. Smith, 2004 Diversity in the Sir2 family of protein deacetylases. J Leukoc Biol 75: 939–950.

Buck, S. W., J. J. Sandmeier and J. S. Smith, 2002 RNA polymerase I propagates unidirectional spreading of rDNA silent chromatin. Cell 111: 1003–1014.

Burton, J. N., I. Liachko, M. J. Dunham and J. Shendure, 2014 Species-level deconvolution of metagenome assemblies with Hi-C-based contact probability maps. G3 (Bethesda) 4: 1339–1346.

Chang, C. R., C. S. Wu, Y. Hom and M. R. Gartenberg, 2005 Targeting of cohesin by transcriptionally silent chromatin. Genes Dev 19: 3031–3042.

Choy, J. S., R. Acuna, W. C. Au and M. A. Basrai, 2011 A role for histone H4K16 hypoacetylation in *Saccharomyces cerevisiae* kinetochore function. Genetics 189: 11–21.

Ciosk, R., M. Shirayama, A. Shevchenko, T. Tanaka, A. Toth et al., 2000 Cohesin’s binding to chromosomes depends on a separate complex consisting of Scc2 and Scc4 proteins. Mol Cell 5: 243–254.

Dang, W., K. K. Steffen, R. Perry, J. A. Dorsey, F. B. Johnson et al., 2009 Histone H4 lysine 16 acetylation regulates cellular lifespan. Nature 459: 802–807.

Dasgupta, A., K. L. Ramsey, J. S. Smith and D. T. Auble, 2004 Sir Antagonist 1 (San1) is a ubiquitin ligase. J Biol Chem 279: 26830–26838.

Defossez, P. A., R. Prusty, M. Kaeberlein, S. J. Lin, P. Ferrigno et al., 1999 Elimination of replication block protein Fob1 extends the life span of yeast mother cells. Mol Cell 3: 447–455.

Derbyshire, M. K., K. G. Weinstock and J. N. Strathern, 1996 *HST1*, a new member of the *SIR2* family of genes. Yeast 12: 631–640.

Duan, Z., M. Andronescu, K. Schutz, S. McIlwain, Y. J. Kim et al., 2010 A three-dimensional model of the yeast genome. Nature 465: 363–367.

Ganley, A. R., and T. Kobayashi, 2014 Ribosomal DNA and cellular senescence: new evidence supporting the connection between rDNA and aging. FEMS Yeast Res 14: 49–59.

Gardner, R. G., Z. W. Nelson and D. E. Gottschling, 2005 Degradation-mediated protein quality control in the nucleus. Cell 120: 803–815.

Gartenberg, M. R., and J. S. Smith, 2016 The nuts and bolts of transcriptionally silent chromatin in *Saccharomyces cerevisiae*. Genetics 203: 1563–1599.

Gomes, A. P., N. L. Price, A. J. Ling, J. J. Moslehi, M. K. Montgomery et al., 2013 Declining NAD+ induces a pseudohypoxic state disrupting nuclear-mitochondrial communication during aging. Cell 155: 1624–1638.

Gotta, M., S. Strahl-Bolsinger, H. Renauld, T. Laroche, B. K. Kennedy et al., 1997 Localization of Sir2p: the nucleolus as a compartment for silent information regulators. EMBO J 16: 3243–3255.

Guacci, V., and D. Koshland, 2012 Cohesin-independent segregation of sister chromatids in budding yeast. Mol Biol Cell 23: 729–739.

Guacci, V., D. Koshland and A. Strunnikov, 1997 A direct link between sister chromatid cohesion and chromosome condensation revealed through the analysis of *MCD1* in *S. cerevisiae*. Cell 91: 47–57.

Heidinger-Pauli, J. M., O. Mert, C. Davenport, V. Guacci and D. Koshland, 2010 Systematic reduction of cohesin differentially affects chromosome segregation, condensation, and DNA repair. Curr Biol 20: 957–963.

Hickman, M. A., and L. N. Rusche, 2007 Substitution as a mechanism for genetic robustness: the duplicated deacetylases Hst1p and Sir2p in *Saccharomyces cerevisiae*. PLoS Genet 3: e126.

Hoggard, T. A., F. Chang, K. R. Perry, S. Subramanian, J. Kenworthy et al., 2018 Yeast heterochromatin regulators Sir2 and Sir3 act directly at euchromatic DNA replication origins. PLoS Genet 14: e1007418.

Hsu, H. C., C. L. Wang, M. Wang, N. Yang, Z. Chen et al., 2013 Structural basis for allosteric stimulation of Sir2 activity by Sir4 binding. Genes Dev 27: 64–73.

Hu, Z., K. Chen, Z. Xia, M. Chavez, S. Pal et al., 2014 Nucleosome loss leads to global transcriptional up-regulation and genomic instability during yeast aging. Genes Dev 28: 396–408.

Huang, J., I. L. Brito, J. Villen, S. P. Gygi, A. Amon et al., 2006 Inhibition of homologous recombination by a cohesin-associated clamp complex recruited to the rDNA recombination enhancer. Genes Dev 20: 2887–2901.

Janssens, G. E., A. C. Meinema, J. Gonzalez, J. C. Wolters, A. Schmidt et al., 2015 Protein biogenesis machinery is a driver of replicative aging in yeast. Elife 4: e08527.

Jessberger, R., 2012 Age-related aneuploidy through cohesion exhaustion. EMBO Rep 13: 539– 546.

Jiang, J. C., E. Jaruga, M. V. Repnevskaya and S. M. Jazwinski, 2000 An intervention resembling caloric restriction prolongs life span and retards aging in yeast. FASEB J 14: 2135–2137.

Kaeberlein, M., M. McVey and L. Guarente, 1999 The SIR2/3/4 complex and SIR2 alone promote longevity in *Saccharomyces cerevisiae* by two different mechanisms. Genes Dev 13: 2570–2580.

Kobayashi, T., and A. R. Ganley, 2005 Recombination regulation by transcription-induced cohesin dissociation in rDNA repeats. Science 309: 1581–1584.

Kobayashi, T., and T. Horiuchi, 1996 A yeast gene product, Fob1 protein, required for both replication fork blocking and recombinational hotspot activities. Genes Cells 1: 465–474.

Kobayashi, T., T. Horiuchi, P. Tongaonkar, L. Vu and M. Nomura, 2004 *SIR2* regulates recombination between different rDNA repeats, but not recombination within individual rRNA genes in yeast. Cell 117: 441–453.

Kogut, I., J. Wang, V. Guacci, R. K. Mistry and P. C. Megee, 2009 The Scc2/Scc4 cohesin loader determines the distribution of cohesin on budding yeast chromosomes. Genes Dev 23: 2345–2357.

Kriegenburg, F., V. Jakopec, E. G. Poulsen, S. V. Nielsen, A. Roguev et al., 2014 A chaperone-assisted degradation pathway targets kinetochore proteins to ensure genome stability. PLoS Genet 10: e1004140.

Kwan, E. X., E. J. Foss, S. Tsuchiyama, G. M. Alvino, L. Kruglyak et al., 2013 A natural polymorphism in rDNA replication origins links origin activation with calorie restriction and lifespan. PLoS Genet 9: e1003329.

Lamming, D. W., M. Latorre-Esteves, O. Medvedik, S. N. Wong, F. A. Tsang et al., 2005 *HST2* mediates *SIR2*-independent life-span extension by calorie restriction. Science 309: 1861– 1864.

Larin, M. L., K. Harding, E. C. Williams, N. Lianga, C. Dore et al., 2015 Competition between heterochromatic loci allows the abundance of the silencing protein, Sir4, to regulate *de novo* assembly of heterochromatin. PLoS Genet 11: e1005425.

Lazar-Stefanita, L., V. F. Scolari, G. Mercy, H. Muller, T. M. Guerin et al., 2017 Cohesins and condensins orchestrate the 4D dynamics of yeast chromosomes during the cell cycle. EMBO J 36: 2684–2697.

Li, C., J. E. Mueller and M. Bryk, 2006 Sir2 represses endogenous polymerase II transcription units in the ribosomal DNA nontranscribed spacer. Mol Biol Cell 17: 3848–3859.

Li, M., B. J. Petteys, J. M. McClure, V. Valsakumar, S. Bekiranov et al., 2010 Thiamine biosynthesis in *Saccharomyces cerevisiae* is regulated by the NAD+-dependent histone deacetylase Hst1. Mol Cell Biol 30: 3329–3341.

Li, M., V. Valsakumar, K. Poorey, S. Bekiranov and J. S. Smith, 2013 Genome-wide analysis of functional sirtuin chromatin targets in yeast. Genome Biol 14: R48.

Lin, S. J., M. Kaeberlein, A. A. Andalis, L. A. Sturtz, P. A. Defossez et al., 2002 Calorie restriction extends *Saccharomyces cerevisiae* lifespan by increasing respiration. Nature 418: 344–348.

Lindstrom, D. L., and D. E. Gottschling, 2009 The mother enrichment program: a genetic system for facile replicative life span analysis in *Saccharomyces cerevisiae*. Genetics 183: 413–422, 411si-413si.

Lindstrom, D. L., C. K. Leverich, K. A. Henderson and D. E. Gottschling, 2011 Replicative age induces mitotic recombination in the ribosomal RNA gene cluster of *Saccharomyces cerevisiae*. PLoS Genet 7: e1002015.

Lister, L. M., A. Kouznetsova, L. A. Hyslop, D. Kalleas, S. L. Pace et al., 2010 Age-related meiotic segregation errors in mammalian oocytes are preceded by depletion of cohesin and Sgo2. Curr Biol 20: 1511–1521.

Lushnikova, T., A. Bouska, J. Odvody, W. D. Dupont and C. M. Eischen, 2011 Aging mice have increased chromosome instability that is exacerbated by elevated Mdm2 expression. Oncogene 30: 4622–4631.

Marston, A. L., 2014 Chromosome segregation in budding yeast: sister chromatid cohesion and related mechanisms. Genetics 196: 31–63.

Matecic, M., D. L. Smith, X. Pan, N. Maqani, S. Bekiranov et al., 2010 A microarray-based genetic screen for yeast chronological aging factors. PLoS Genet 6: e1000921.

Matheson, T. D., and P. D. Kaufman, 2016 Grabbing the genome by the NADs. Chromosoma 125: 361–371.

Mehta, R., D. Chandler-Brown, F. J. Ramos, L. S. Shamieh and M. Kaeberlein, 2010 Regulation of mRNA translation as a conserved mechanism of longevity control. Adv Exp Med Biol 694: 14–29.

Moazed, D., A. Kistler, A. Axelrod, J. Rine and A. D. Johnson, 1997 Silent information regulator protein complexes in *Saccharomyces cerevisiae*: a SIR2/SIR4 complex and evidence for a regulatory domain in SIR4 that inhibits its interaction with SIR3. Proc Natl Acad Sci U S A 94: 2186–2191.

Mortimer, R. K., and J. R. Johnston, 1959 Life span of individual yeast cells. Nature 183: 1751–1752.

Nagaoka, S. I., T. J. Hassold and P. A. Hunt, 2012 Human aneuploidy: mechanisms and new insights into an age-old problem. Nat Rev Genet 13: 493–504.

Neurohr, G. E., R. L. Terry, A. Sandikci, K. Zou, H. Li et al., 2018 Deregulation of the G1/S-phase transition is the proximal cause of mortality in old yeast mother cells. Genes Dev 32: 1075–1084.

North, B. J., M. A. Rosenberg, K. B. Jeganathan, A. V. Hafner, S. Michan et al., 2014 SIRT2 induces the checkpoint kinase BubR1 to increase lifespan. Embo j 33: 1438–1453.

O’Sullivan, J. M., D. M. Sontam, R. Grierson and B. Jones, 2009 Repeated elements coordinate the spatial organization of the yeast genome. Yeast 26: 125–138.

Ohkuni, K., and K. Kitagawa, 2011 Endogenous transcription at the centromere facilitates centromere activity in budding yeast. Curr Biol 21: 1695–1703.

Oppikofer, M., S. Kueng, F. Martino, S. Soeroes, S. M. Hancock et al., 2011 A dual role of H4K16 acetylation in the establishment of yeast silent chromatin. EMBO J 30: 2610–2621.

Pal, S., S. D. Postnikoff, M. Chavez and J. K. Tyler, 2018 Impaired cohesion and homologous recombination during replicative aging in budding yeast. Sci Adv 4: eaaq0236.

Perkins, A. T., T. M. Das, L. C. Panzera and S. E. Bickel, 2016 Oxidative stress in oocytes during midprophase induces premature loss of cohesion and chromosome segregation errors. Proc Natl Acad Sci U S A 113: E6823–e6830.

Riesen, M., and A. Morgan, 2009 Calorie restriction reduces rDNA recombination independently of rDNA silencing. Aging Cell 8: 624–632.

Rine, J., and I. Herskowitz, 1987 Four genes responsible for a position effect on expression from *HML* and *HMR* in *Saccharomyces cerevisiae*. Genetics 116: 9–22.

Sasaki, T., B. Maier, A. Bartke and H. Scrable, 2006 Progressive loss of SIRT1 with cell cycle withdrawal. Aging Cell 5: 413–422.

Schalbetter, S. A., A. Goloborodko, G. Fudenberg, J. M. Belton, C. Miles et al., 2017 SMC complexes differentially compact mitotic chromosomes according to genomic context. Nat Cell Biol 19: 1071–1080.

Schlissel, G., M. K. Krzyzanowski, F. Caudron, Y. Barral and J. Rine, 2017 Aggregation of the Whi3 protein, not loss of heterochromatin, causes sterility in old yeast cells. Science 355: 1184–1187.

Shou, W., J. H. Seol, A. Shevchenko, C. Baskerville, D. Moazed et al., 1999 Exit from mitosis is triggered by Tem1-dependent release of the protein phosphatase Cdc14 from nucleolar RENT complex. Cell 97: 233–244.

Sinclair, D. A., and L. Guarente, 1997 Extrachromosomal rDNA circles--a cause of aging in yeast. Cell 91: 1033–1042.

Smeal, T., J. Claus, B. Kennedy, F. Cole and L. Guarente, 1996 Loss of transcriptional silencing causes sterility in old mother cells of *S. cerevisiae*. Cell 84: 633–642.

Smith, D. L., Jr., C. Li, M. Matecic, N. Maqani, M. Bryk et al., 2009 Calorie restriction effects on silencing and recombination at the yeast rDNA. Aging Cell 8: 633–642.

Smith, J. S., and J. D. Boeke, 1997 An unusual form of transcriptional silencing in yeast ribosomal DNA. Genes Dev 11: 241–254.

Smith, J. S., C. B. Brachmann, L. Pillus and J. D. Boeke, 1998 Distribution of a limited Sir2 protein pool regulates the strength of yeast rDNA silencing and is modulated by Sir4p. Genetics 149: 1205–1219.

Smith, J. S., E. Caputo and J. D. Boeke, 1999 A genetic screen for ribosomal DNA silencing defects identifies multiple DNA replication and chromatin-modulating factors. Mol Cell Biol 19: 3184–3197.

Song, S., and F. B. Johnson, 2018 Epigenetic mechanisms impacting aging: a focus on histone levels and telomeres. Genes (Basel) 9: 201.

Spencer, F., S. L. Gerring, C. Connelly and P. Hieter, 1990 Mitotic chromosome transmission fidelity mutants in *Saccharomyces cerevisiae*. Genetics 124: 237–249.

Steffen, K. K., B. K. Kennedy and M. Kaeberlein, 2009 Measuring replicative life span in the budding yeast. J Vis Exp. 28: e1209.

Straight, A. F., W. Shou, G. J. Dowd, C. W. Turck, R. J. Deshaies et al., 1999 Net1, a Sir2-associated nucleolar protein required for rDNA silencing and nucleolar integrity. Cell 97: 245–256.

Sunshine, A. B., G. T. Ong, D. P. Nickerson, D. Carr, C. J. Murakami et al., 2016 Aneuploidy shortens replicative lifespan in *Saccharomyces cerevisiae*. Aging Cell 15: 317–324.

Takeuchi, Y., T. Horiuchi and T. Kobayashi, 2003 Transcription-dependent recombination and the role of fork collision in yeast rDNA. Genes Dev 17: 1497–1506.

Uhlmann, F., 2016 SMC complexes: from DNA to chromosomes. Nat Rev Mol Cell Biol 17: 399–412.

Unal, E., J. M. Heidinger-Pauli, W. Kim, V. Guacci, I. Onn et al., 2008 A molecular determinant for the establishment of sister chromatid cohesion. Science 321: 566–569.

Utani, K., and M. I. Aladjem, 2018 Extra View: Sirt1 Acts as a gatekeeper of replication initiation to preserve genomic stability. Nucleus 9: 261–267.

Wasko, B. M., and M. Kaeberlein, 2014 Yeast replicative aging: a paradigm for defining conserved longevity interventions. FEMS Yeast Res 14: 148–159.

Winston, F., C. Dollard and S. L. Ricupero-Hovasse, 1995 Construction of a set of convenient *Saccharomyces cerevisiae* strains that are isogenic to S288C. Yeast 11: 53–55.

Wood, A. J., A. F. Severson and B. J. Meyer, 2010 Condensin and cohesin complexity: the expanding repertoire of functions. Nat Rev Genet 11: 391–404.

Wu, C. S., Y. F. Chen and M. R. Gartenberg, 2011 Targeted sister chromatid cohesion by Sir2. PLoS Genet 7: e1002000.

Yoshida, K., J. Bacal, D. Desmarais, I. Padioleau, O. Tsaponina et al., 2014 The histone deacetylases Sir2 and Rpd3 act on ribosomal DNA to control the replication program in budding yeast. Mol Cell 54: 691–697.

Zhu, J., D. Heinecke, W. A. Mulla, W. D. Bradford, B. Rubinstein et al., 2015 Single-cell based quantitative assay of chromosome transmission fidelity. G3 (Bethesda) 5: 1043–1056.

