## Supplemental Information for "Depletion of limiting rDNA structural complexes 1 triggers chromosomal instability and replicative aging of *Saccharomyces cerevisiae*"

### Supplemental Information (Fine et al.)

#### Table S1. Yeast Strains

| Strain | Genotype | Source |
|---|---|---|
| BY4741 | *MAT*a *his3Δ1 leu2Δ0 LYS2 met15Δ0 ura3Δ0* | (BRACHMANN *et al.* 1998) |
| BY4742 | *MATα his3Δ*1 *leu2Δ0 lys2Δ0 MET15 ura3Δ0* | (BRACHMANN *et al.* 1998) |
| FY834 | *MATα leu2Δ1 lys2Δ202 trp1Δ63 his3Δ200 ura3*-52 | (Winston *et al.* 1995) |
| ML1 | *MAT*a *his3Δ200 leu2Δ1 met15Δ0 trp1Δ63 ura3-167* | (LI *et al.* 2010) |
| ML150 | *MAT*a *his3Δ200 leu2Δ1 met15Δ0 trp1Δ63 ura3-167 MCD1-13xMyc:kanMX4* | (LI *et al.* 2010) |
| UCC5179 | *MAT*a *ade2::hisG his3 leu2 lys2 ura3Δ0 trpΔ63 HOΔ::SCW11 pr-Cre-EBD78-natMX loxP-UBC9-loxP-LEU2 loxP-CDC20-loxP-hphMX* | (LINDSTROM AND GOTTSCHLING 2009) |
| UCC5181 | *MATα ade2::hisG his3 leu2 met15Δ::ADE2 ura3Δ0 trpΔ63 HOΔ::SCW11 pr-Cre-EBD78-natMX loxP-UBC9-loxP-LEU2 loxP-CDC20-loxP-hphMX* | (LINDSTROM AND GOTTSCHLING 2009) |
| UCC5185 | UCC5179/UCC5181 | (LINDSTROM AND GOTTSCHLING 2009) |
| YPH278 | *MATα ura3-52 lys2-801 ade2-101 his3Δ200 leu2Δ1 CFIII [pCEN3.L.YPH278, SUP11, URA3]* | (SPENCER *et al.* 1990) |
| RF3 | UCC5179*; MCD1-13xMyc-kanMX4* | This Study |
| RF4 | UCC5181*;MCD1-13xMyc-kanMX4* | This Study |
| RF10 | RF3/RF4*; MCD1-13xMyc-kanMX4/MCD1-13xMyc-kanMX4* | This Study |
| RF30 | UCC5181*; SCC2-13xMyc-kanMX4* | This Study |
| RF31 | UCC5179*;SCC2-13xMyc-kanMX4* | This Study |
| RF32 | YPH278*; sir2Δ::natMX4* | This Study |
| RF33 | YPH278*; hst1Δ::natMX4* | This Study |
| RF34 | RF30/RF31*; SCC2-13xMyc-kanMX4/SCC2-13xMyc-kanMX4* | This Study |
| 3349-1B | *LacO [DK] NAT::lys4 trp1-1 bar1 [pHIS3, GFP-LacI, HIS3]::his3-11,15 leu2-3,112 ura3-52 GAL*$^{+}$ | (GUACCI *et al.* 1997) |

| | | |
|---|---|---|
| 3312-7A | *mcd1-1 LacO-NAT::lys4 trp1-1 bar1 [pHIS3,GFP-LacI, HIS3]::his3-11,15 leu2-3,112 ura2-52 GAL*$^+$ | (GUACCI *et al.* 1997) |
| 3460-2A | *LacO [DK]-NAT: 10kb CENIV trp1-1 bar1 [pHIS3, GFPLacI, HIS3]::his3-11,15 leu2-3,112 ura2-52 GAL*$^+$ | (GUACCI *et al.* 1997) |
| RF43 | YPH278*; hst1Δ::natMX4, sir2Δ::kanMX4* | This Study |
| RF77 | UCC5181*; SMC1-13xMyc-kanMX4* | This Study |
| JH5275b | *MATalphaΔ::hphMX hoΔ hml1Δ::ADE1 hmrΔ::ADE1 ade1-110 leu2,3-112::MCD1-LEU2 lys5 trp1::hisG ura3-52 ade3::GAL10:HO met17::trna-SUP53-G418 mcd1Δ::natMX* | (HEIDINGER-PAULI *et al.* 2010) |
| JH5276b | *MATalphaΔ::hphMX hoΔ hml1Δ::ADE1 hmrΔ::ADE1 ade1-110 leu2,3-112::mcd1L12STOP-LEU2 lys5 trp1::hisG ura3-52 ade3::GAL10:HO met17::trna-SUP53-G418 mcd1Δ::natMX* | (HEIDINGER-PAULI *et al.* 2010) |
| RF89 | UCC5179*;SMC1-13xMyc-kanMX4* | This Study |
| RF90 | RF77/RF89*; SMC1-13xMyc-kanMX4/SMC1-13xMyc-kanMX4* | This Study |
| RF113 | JH5275b*; ura3-52::[pRS306, Empty, URA3]* | This Study |
| RF114 | JH5276b*; ura3-52::[pRS306, Empty, URA3]* | This Study |
| RF118 | JH5275b*; ura3-52::[pJSB186, SIR2, URA3]* | This Study |
| RF119 | JH5276b*; ura3-52::[pJSB186, SIR2, URA3]* | This Study |
| RF127 | YPH278*; sir2Δ::kanMX4* | This Study |
| RF131 | YPH278*; hst1Δ::natMX4, sir2Δ::kanMX4 fob1Δ::hphMX4* | This Study |
| RF132 | YPH278*; sir2Δ::kanMX4 hmrΔ::hphMX4* | This Study |
| RF134 | YPH278*;hst1Δ::natMX4, sir2Δ::kanMX4 hmrΔ::hphMX4* | This Study |
| RF135 | YPH278*;hst1Δ::natMX4, hmrΔ::hphMX4* | This Study |
| RF137 | YPH278*; hmrΔ::hphMX4* | This Study |
| RF138 | YPH278*; hst1Δ::natMX4, fob1Δ::hphMX4* | This Study |
| RF146 | JH5276B*; [pCM252, pTET::Empty, TRP1]* | This Study |
| RF147 | JH5275B*; [pCM252, pTET::Empty, TRP1]* | This Study |
| RF148 | YPH278*; fob1Δ::hphMX4* | This Study |
| RF149 | YPH278*; fob1Δ::hphMX4 sir2Δ::kanMX4* | This Study |
| RF176 | UCC5181*; LRS4-13xMyc::kanMX4* | This Study |
| RF179 | JH5275B*; [pRF4, pTET::MCD1, TRP1]* | This Study |
| RF180 | JH5276B*; [pRF4, pTET::MCD1, TRP1]* | This Study |
| RF190 | UCC5181*; SIR4-13xMyc-kanMX4* | This Study |
| RF204 | UCC5181*; MATa NET1-13xMyc::kanMX4 met15*[1] | This Study |
| RF206 | UCC5179*; LRS4-13xMyc-kanMX4* | This Study |
| RF207 | RF196/RF206*; LRS4-13xMyc-kanMX4/LRS4-13xMyc-kanMX4* | This Study |
| W303AR | *RDN::ADE2 leu2-3,112 trp1-1 can1-100 ura3-1 ade2-1 his3-11,15* | (KAEBERLEIN *et al.* 1999) |
| DSY1034 | *RDN::ADE2 leu2-3,112 trp1-1 can1-100 ura3-1 ade2-1 his3-11,15 sir2Δ::TRP1* | (KAEBERLEIN *et al.* 1999) |
| RF212 | UCC5181; *NET1-13xMyc-kanMX4 met15::ADE2* | This Study |
| RF213 | RF204/RF212*; NET1-13xMyc-kanMX4/NET1-13xMyc-kanMX4 met15::ADE2/met15*[1] | This Study |

| | | |
|---|---|---|
| RF217 | YPH278; *MAT*a *fob1Δ::hphMX4 sir2Δ::kanMX* | This Study |
| RF218 | YPH278; *leu2Δ1::[pRF10, pTET::Empty, LEU2]* | This Study |
| RF219 | YPH278; *leu2Δ1::[pRF11, pTET::MCD1, LEU2]* | This Study |
| RF222 | YPH278; *sir2Δ::kanMX leu2Δ1::[pRF10, pTET::Empty, LEU2]* | This Study |
| RF223 | YPH278; *sir2Δ::kanMX leu2Δ1::[pRF11, pTET::MCD1, LEU2]* | This Study |
| RF224 | YPH278; *sir2Δ::kanMX hst1Δ::natMX4 leu2Δ1::[pRF10, pTET::Empty, LEU2]* | This Study |
| RF225 | YPH278; *sir2Δ::kanMX hst1Δ::natMX4 leu2Δ1::[pRF11, pTET::MCD1, LEU2]* | This Study |
| RF226 | W303AR; *leu2Δ1::[pRF10, pTET::Empty, LEU2]* | This Study |
| RF227 | W303AR; *leu2Δ1::[pRF11, pTET::MCD1, LEU2]* | This Study |
| RF228 | DSY1034; *[pRF10, pTET::Empty, LEU2]::leu2Δ1* | This Study |
| RF229 | DSY1034; *[pRF11, pTET::MCD1, LEU2]::leu2Δ1* | This Study |
| RF232 | YPH278; *sir2Δ::kanMX hst1Δ::natMX4 hmrΔ::hphMX fob1Δ::hphMX* | This Study |
| RF242 | UCC5179; *SIR4-13xMyc::kanMX4, lys2*[1] *met15::ADE2* | This Study |
| RF243 | RF190/RF242; *SIR4-13xMyc::kanMX4/ SIR4-13xMyc::kanMX4 lys2*[1]*/LYS2 met15::ADE2/met15::ADE2* | This Study |
| RF258 | 3460-2A; *SPB1-dsRED-kanMX4* | This Study |
| RF278 | 3460-2A; *SPB1-dsRED-kanMX4 fob1Δ::hphMX4* | |
| RF288 | FY834; *[pRF11, pTET::MCD1, LEU2]::leu2Δ1* | This Study |
| RF289 | FY834; *[pRF10, pTET::Empty, LEU2]::leu2Δ1* | This Study |
| RF290 | 3460-2A; *SPB1-dsRED-kanMX4 sir2Δ::hphMX4* | |

[1]Allele status unknown

**Table S2. Oligonucleotides used in the study.**

| Primer | Target | Sequence |
|---|---|---|
| JS369 | *fob1Δ*_Det_2793 | GACGACAATACCGCTGGCTCCCGT |
| JS646 | *SIR4*-Myc_Fw | GGAAAAAGATTTTCAAGTGAATAAGGAGATAAAACCGTATCGGATCCCCGGGTTAATTAA |
| JS647 | *SIR4*-Myc_Rv | GGTACACTTCGTTACTGGTCTTTTGTAGAATGATAAAAAGGAATTCGAGCTCGTTTAAAC |
| JS900 | *fob1Δ*_hphMX_Fw | GGAGAACAATTTAACGATTGTGTGAGTAATTTGTGCTCCGGCGCGAAGCAAAAATTACGGC |
| JS901 | *fob1Δ*_hphMX_Rv | CACCTATGACTCCTCCTTTCATTCTATCCTACATATTCGGCGTTAGTATCGAATCGACAGC |
| JS1100 | NTS1_Fw | TGTTAGTGCAGGAAAGCGGG |
| JS1101 | NTS1_Rv | CTACACCCTCGTTTAGTTGC |
| JS1146 | *ACT1* RT-qPCR_Fw | ATGTGTAAAGCCGGTTTTGCC |
| JS1147 | *ACT1* RT-qPCR_Rv | TGGGAAGACAGCACGAGGAG |
| JS2129 | *MCD1*-Myc_Fw | AGACGCCAAACCTGCACTATTTGAAAGGTTTATCAATGCTCGGATCCCCGGGTTAATTAA |
| JS2130 | *MCD1*-Myc_Rv | TATTGGGTCCACCAAGAAATCCCCTCGGCGTAACTAGGTTGAATTCGAGCTCGTTTAAAC |
| JS2131 | *SCC2*-Myc_Fw | CAAGCTTCTTACATATTTTAGAAAACACGTGAAGGATACGCGGATCCCCGGGTTAATTAA |
| JS2132 | *SCC2*-Myc_Rv | AATGATTATTAATACTATGTATATTTTAAGTGCAATATATGAATTCGAGCTCGTTTAAAC |
| JS2153 | *sir2Δ*_natMX_Fw | CATTCAAACCATTTTTCCCTCATCGGCACATTAAAGCTGGCCGGCGCGAAGCAAAAATTACGGC |
| JS2154 | *sir2Δ*_natMX_Rv | TATTAATTTGGCACTTTTAAATTATTAAATTGCCTTCTACCGGCGTTAGTATCGAATCGACAGC |
| JS2155 | *hst1Δ*_natMX_Fw | CTCTTCTTTTTTGTTGTTTTGTGAGAAAAAAAAATCTAA CCGGCGCGAAGCAAAAATTACGGC |
| JS2156 | *hst1Δ*_natMX_Rv | TGCAATAGCAGCGGTATACTTATTTTTACTCCCCCTTCTG CGGCGTTAGTATCGAATCGACAGC |
| JS2167 | *PDC1*_Fw | GCCGACAGTCTGTTGAATTGG |
| JS2168 | *PDC1*_Rv | GAAGCGGACCCAGACTTAAGC |
| JS2300 | *TELXV*_Rv | TGGGTAAATGGCAAAGGGTA |
| JS2301 | *TELXV*_Fw | GCACCCACATCATTATGCAC |
| JS2478 | *natMX*_C_Readout | CCTGGACACCGCCCTGTA |
| JS2480 | *hst1Δ*_Det_JS2478 | CGGACATTTTCATTCAAGAGG |
| JS2602 | *CEN3*_Fw | AGCGCCAAACAATATGGAAA |
| Js2603 | *CEN3*_Rv | TGAGCAAAACTTCCACCAGT |
| JS2606 | 10kb right of *CEN3*_Fw | CCTGCGTCACACATGAGAAA |
| JS2607 | 10kb right of *CEN3*_Rv | TCACAGTTTACCCGGAGGTC |
| JS2608 | *CEN4*_Fw | ATTGCTTGCAAAAGGTCACA |
| JS2609 | *CEN4*_Rv | GCCGCTCCTAGGTAGTGCTT |

| | | |
|---|---|---|
| JS2734 | *SMC1*-Myc_Fw | GTCGAAGATCATAACTTTGGACTTGAGCAATTACGCAG AACGGATCCCCGGGTTAATTAA |
| JS2735 | *SMC1*-Myc_Rv | TAGTTATTTGACGGGTTATAGCAGAGGTTGGTTTCATAG AGAATTCGAGCTCGTTTAAC |
| JS2757 | *CEN11*_Fw | CCTAATACCTCAATGGTCCAATAC |
| JS2758 | *CEN11*_Rv | TGCTTTGATTTATACAGAGGAAGC |
| JS2763 | *STE2*_Fw | TTCGTGACCTTCGGTATAAGG |
| JS2764 | *STE2*_Rv | CGTAAAAGCAAAGGTGGTTTCT |
| JS2789 | *LRS4*-Myc_Fw | TGAAGAATTGAGTAATAATTTAAATGTTGACGAGTTTGTA CGGATCCCCGGGTTAATTAA |
| JS2790 | *LRS4*-Myc_Rv | AAAAGAGAGAGGGAAGGGCAGGGACGTAGATAGCTGT TACGAATTCGAGCTCGTTTAAAC |
| JS2793 | *hphMX*_N_Readout | CTTCCGGAATCGGGAGCGCG |
| JS2813 | *hmrΔ*_hphMX_Fw | CGGGCTCATTCTTTCTTCTTTGCCAGAGGCTCACCGCCG GCGCGAAGCAAAAATTACGGC |
| JS2814 | *hmrΔ*_hphMX_Rv | ACGAATTTATTTAGATCTCATACGTTTATTTATGAACGG CGTTAGTATCGAATCGACAGC |
| JS2819 | *hmrΔ*_Det_JS2793 | TGAGAATAAGCGCAGGTACTCCTGGT |
| JS2864 | *MCD1*_ORF_*Not*I _Fw | GAATTTGCGGCCGCATGGTTACAGAAAATCCTCAACGT CTTAC |
| JS2865 | *MCD*1_ORF_*Pst*I_Rv | GAATTTCTGCAGCGGTAGAGGTGTGGTCAATAAGAG |
| JS2842 | 15kb right of *CEN4*_Fw | TCAGTGCATCCATCGTCTTC |
| JS2843 | 15kb right of *CEN4*_Rv | TTGAGCTTCTAGCGACAGCA |
| JS2844 | *MCD1* RT-qPCR_Fw | ACAGAAAGAGAAGTTTTGGTCACC |
| JS2949 | *MCD1* RT-qPCR_Rv | TGAAGTATCCCAGGGAGCAG |
| JS3028 | 200kb right of *CEN4*_Fw | TTTTGGCACAAAGGCAATG |
| JS3029 | 200kb right of *CEN4*_Rv | TGAATCCATTTGTTTGTTAATAGTTT |
| JS3034 | *CEN6*_Fw | TTGAAGACTATATTTCTTTTCATCACG |
| JS3035 | *CEN6*_Rv | AACTTTCAACCTATTTTACATCTTCG |
| JS3036 | *CEN14*_Fw | GCAATTAATTGAGTTGTTGTGAAATGG |
| JS3037 | *CEN14*_Rv | CGCAGATATCCTTAAATACCAAA |

**Table S3. Detailed bud scar and western blot quantitation data.**

| Microscopy | | | Western Blot | |
|---|---|---|---|---|
| Factor | Bud Scars | St. Error | Qty. | St. Error |
| Mcd1_young | 0.833333 | 0.272845 | 1 | 0 |
| Mcd1_old | 6.400877 | 0.445052 | 0.146018 | 0.119223 |
| Mcd1_CR_young | 0.216667 | 0.044096 | 1 | 0 |
| Mcd1_CR_old | 4.793333 | 0.716038 | 0.252445 | 0.08759 |
| Sir2_young | 0.975362 | 0.114328 | 1 | 0 |
| Sir2_old | 5.573099 | 0.777897 | 0.228414 | 0.186502 |
| Scc2_young | 0.54386 | 0.072505 | 1 | 0 |
| Scc2_old | 6.416667 | 0.469338 | 0.436795 | 0.083468 |
| Smc1_young | 0.637009 | 0.235205 | 1 | 0 |
| Smc1_old | 4.951629 | 0.226633 | 0.188713 | 0.154083 |
| Hst1_young | 0.508696 | 0.158696 | 1 | 0 |
| Hst1_old | 4.919553 | 0.450378 | 0.37225 | 0.111099 |
| Net1_young | 0.25 | 0.028868 | 1 | 0 |
| Net1_old | 5.328571 | 0.289469 | 0 | 0 |
| Lrs4_young | 0.066667 | 0.033333 | 1 | 0 |
| Lrs4_old | 5.049206 | 0.262414 | 0.077961 | 0.063655 |
| Sir3_young | 0.483333 | 0.208833 | 1 | 0 |
| Sir3_old | 4.995238 | 0.145238 | 1.229025 | 0.244495 |
| Sir4_young | 0.825 | 0.375 | 1 | 0 |
| Sir4_old | 6.0375 | 0.9875 | 0.138803 | 0.098148 |

**Supplemental References**

Brachmann, C. B., A. Davies, G. J. Cost, E. Caputo, J. Li *et al.*, 1998 Designer deletion strains derived from *Saccharomyces cerevisiae* S288C: a useful set of strains and plasmids for PCR-mediated gene disruption and other applications. Yeast 14**:** 115-132.

Guacci, V., D. Koshland and A. Strunnikov, 1997 A direct link between sister chromatid cohesion and chromosome condensation revealed through the analysis of MCD1 in *S. cerevisiae.* Cell 91**:** 47-57.

Heidinger-Pauli, J. M., O. Mert, C. Davenport, V. Guacci and D. Koshland, 2010 Systematic reduction of cohesin differentially affects chromosome segregation, condensation, and DNA repair. Curr Biol 20**:** 957-963.

Kaeberlein, M., M. McVey and L. Guarente, 1999 The SIR2/3/4 complex and SIR2 alone promote longevity in *Saccharomyces cerevisia*e by two different mechanisms. Genes Dev 13**:** 2570-2580.

Li, M., B. J. Petteys, J. M. McClure, V. Valsakumar, S. Bekiranov *et al.*, 2010 Thiamine biosynthesis in *Saccharomyces cerevisiae* is regulated by the $NAD^+$-dependent histone deacetylase Hst1. Mol Cell Biol 30**:** 3329-3341.

Lindstrom, D. L., and D. E. Gottschling, 2009 The mother enrichment program: a genetic system for facile replicative life span analysis in *Saccharomyces cerevisiae*. Genetics 183**:** 413-422, 411si-413si.

Spencer, F., S. L. Gerring, C. Connelly and P. Hieter, 1990 Mitotic chromosome transmission fidelity mutants in *Saccharomyces cerevisiae*. Genetics 124**:** 237-249.

Winston, F., C. Dollard, and S.L. Ricupero-Hovasse, 1995 Construction of a set of convenient *Saccharomyces cerevisiae* strains that are isogenic to S288C. Yeast 11: 53-55.

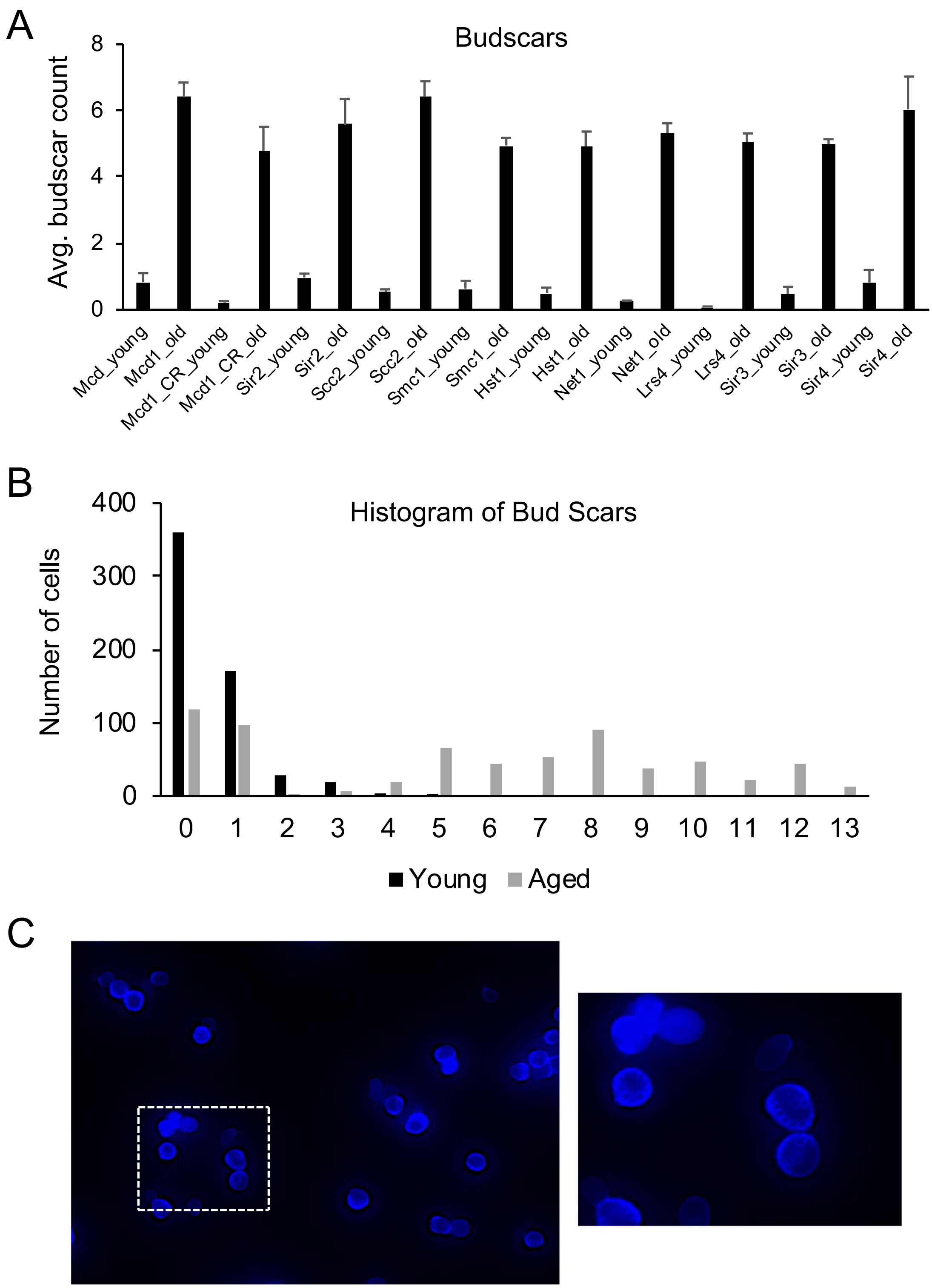
A
Budscars
Avg. budscar count
8
6
4
2
0
Mcd_young
Mcd1_old
Mcd1_CR_young
Mcd1_CR_old
Sir2_young
Sir2_old
Scc2_young
Scc2_old
Smc1_young
Smc1_old
Hst1_young
Hst1_old
Net1_young
Net1_old
Lrs4_young
Lrs4_old
Sir3_young
Sir3_old
Sir4_young
Sir4_old
B
Histogram of Bud Scars
Number of cells
400
300
200
100
0
0 1 2 3 4 5 6 7 8 9 10 11 12 13
Young
Aged
C

**Figure S1. Detailed western blot and bud scar count quantification. (A)** Bar graph showing average bud scars from young and aged cell populations that were used for western blotting. **(B)** Histogram of bud scars from young and aged populations used in western blot experiments. (n=584 and 658) **(C)** Representative image of an enriched aged cell population from the third Lrs4-13xMyc biological replicate. Inset image allows counting of individual bud scars.

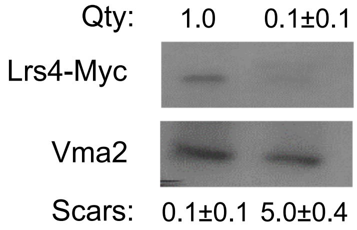


**Figure S2. The Lrs4 subunit of cohibin/monopolin is depleted in aged yeast cells.** Representative western blot of 13xMyc-tagged Lrs4. Vma2 is used as a loading control. Average bud scar counts are indicated at the bottom, and relative Lrs4-myc signal compared to the Vma2 loading control is indicated at the top (Qty).

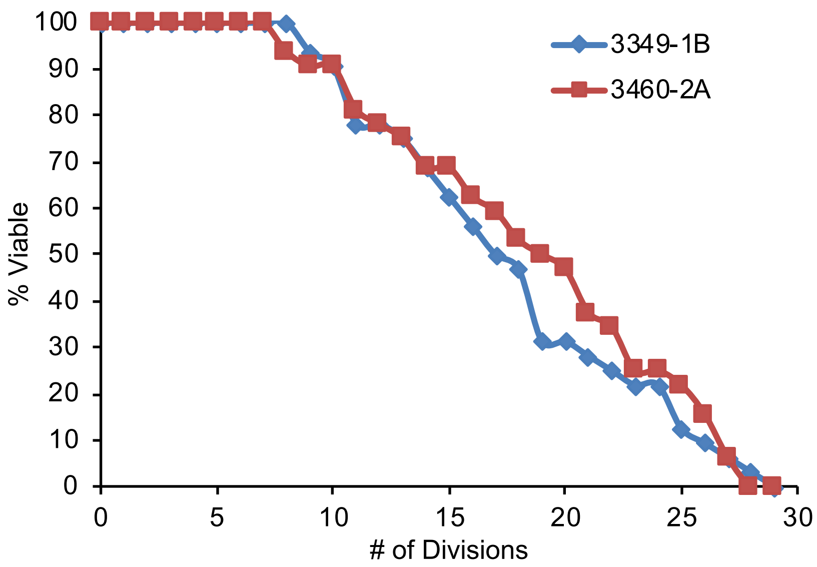


**Figure S3. RLS of the cohesion visualization strains is normal and unaffected by position of the lacO array.** Strain 3349-1B contains a lacO array at the *LYS4* locus on ChrIV and is used as a proxy for arm cohesion, while strain 3460-2A contains a lacO array on ChrIV 10 kb away from the *CEN4* locus that is used to monitor centromeric cohesion.

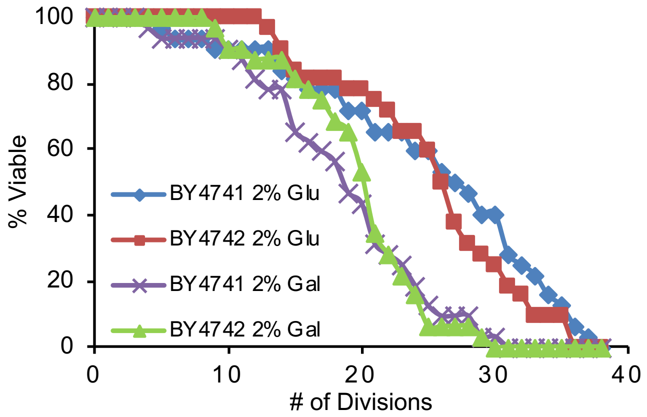


**Figure S4. Galactose shortens yeast RLS in the common yeast BY4741/4742 strain background. (A)** RLS of WT BY4741(*MAT*a) and BY4742 (*MAT*a) cells (n=32; mean rls:◆24.3, ■24.2, ▲17.7, ×18.9) growing on YEP agar plates containing either 2% glucose or galactose as the carbon source.

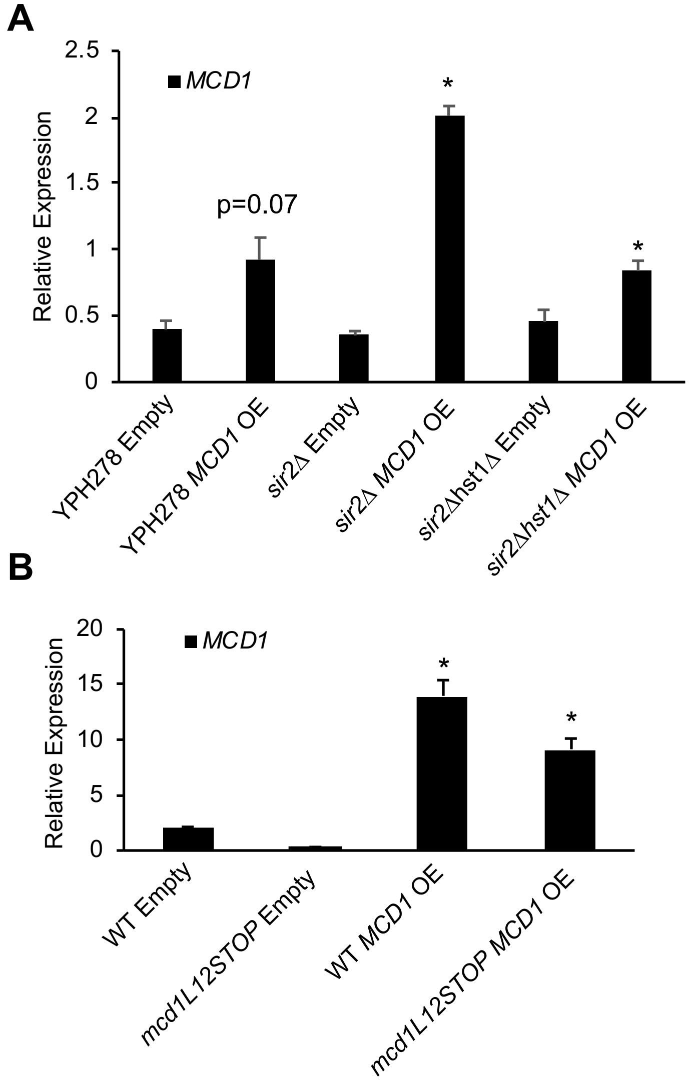


**Figure S5. Doxycycline-induced *MCD1* overexpression in WT and *mcd1L12STO*P strains. (A)** RT-qPCR of *MCD1* transcript levels relative to actin transcript levels in chromosome loss assay strains (YPH background) used to examine *SIR2* and *MCD1* epistasis in RLS. **(B)** RT-qPCR of *MCD1* transcript levels relative to actin transcript levels were quantified from the empty vector strains (RF146 and RF147) or the *MCD1* overexpression strains RF179 and RF180. Total RNA was isolated following 4 hours doxycycline induction during log-phase growth. (*$p<0.05$, student's two-tailed t-test).

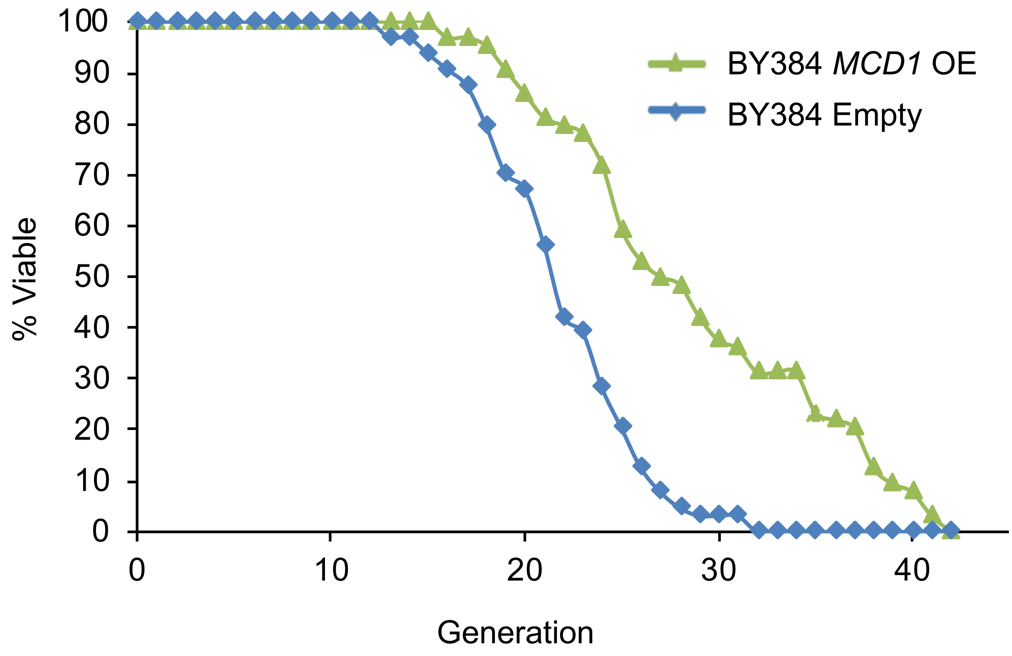


**Figure S6. Doxycycline-induced *MCD1* overexpression in BY Background.** Replicative lifespan viability assay in which an integrated inducible tetracycline promoter overexpressed either the *MCD1* gene or an empty vector (mean RLS = ▲ 28.0 ♦ 21.0).

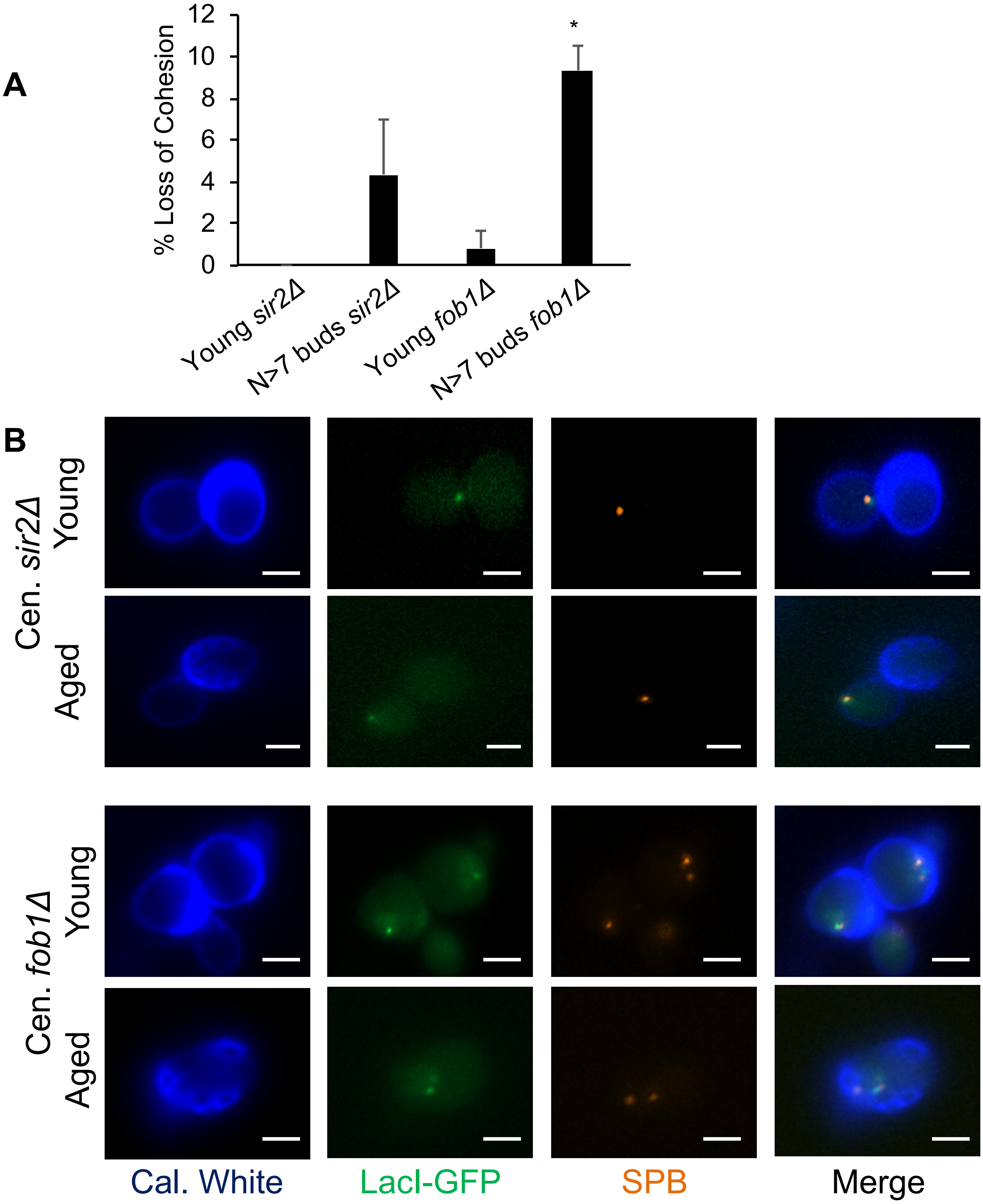


**Figure S7. rDNA stability does not effect sister chromatid cohesion. (A)** Quantification of cells assayed in **(B)** for loss (2-GFP Dots) of chromatid cohesion. (n=60 cells) **(B)** Representative screen shots of young (log-phase) or aged yeast cells monitoring sister chromatid cohesion 10kb proximal to *CENIV* in *sir2Δ* and *fob1Δ* mutants.
